# A platform for robust quantitative functional genomics reveals temporal fitness landscapes in African trypanosomes

**DOI:** 10.64898/2026.06.27.734978

**Authors:** Simon D’Archivio, Lorenzo Brusini, Sandra Trindade, Luisa M Figueiredo, Catarina Gadelha, Bill Wickstead

**Author notes:** these authors contributed equally to this work (Simon D’Archivio, Lorenzo Brusini).

## Abstract

Genome-scale phenotypic screening is transforming our understanding of host-pathogen interactions, but accurate quantitation of mutant fitness and generation of libraries in strains that capture complex disease characteristics remains challenging. Here we introduce Direct RNAi Fragment Sequencing (DRiF-Seq), a sequencing-optimized high-throughput RNA-interference platform that enables robust, clone-resolved quantification of fitness effects in African trypanosomes. Combined with an a posteriori noise estimation approach, DRiF-Seq provides reproducible estimates of fitness magnitude, timing, and statistical confidence for individual mutants and genes. Application to pleomorphic bloodstream-form *Trypanosoma brucei* enables the first genome-scale quantitative profiling in a mammalian infection model, creating a framework for investigating parasite biology at scale in host. DRiF-Seq demonstrates that over half of core genes have a fitness cost when targetted by RNAi, accurately resolves fitness landscapes within molecular complexes, identifies characteristic temporal signatures of gene disruption, and detects biologically meaningful differences between closely related parasite strains that predict differential drug sensitivity. Comparative analyses with genome-wide datasets from *Toxoplasma* and *Plasmodium* further demonstrate conserved and lineage-specific determinants of parasite fitness across eukaryotes. Together, DRiF-Seq provides a sensitive and transferable framework for quantitative functional genomics, a community resource for understanding parasite biology and a system for exploitation of high-complexity mutant screens in host.

## Introduction

Understanding gene function is fundamental to decoding the complexities of biological systems, elucidating disease mechanisms, and identifying the modes of action for, and mechanisms of resistance to, therapeutic interventions. High-throughput phenotypic screens have emerged as transformative tools in the field of functional genomics, enabling researchers to systematically perturb and analyse the roles of genes across the entire genome. These screens are particularly powerful due to their ability to interrogate a vast number of mutants simultaneously and quantitatively – and have the potential to provide robust measurements of gene perturbation effects that afford comprehensive insights into genetic networks.

The application of high-throughput phenotypic screens in protozoan parasites is revolutionizing our approach to investigating the biology of important pathogens. In *Toxoplasma gondii*, genome-wide measurement of growth of mutants induced by CRISPR/Cas9 identified previously unknown genes critical for parasite survival (Sidik et al. 2016) in addition to genes involved in parasite response to drugs (Harding et al. 2020), oxidative stress (Chen et al. 2021), and interferon gamma-mediated control (Wang et al. 2020). In *Plasmodium berghei* a different approach using vector insertion has been used to individually knockout 2,578 genes (∼50% of the protein-coding genes) to systematically measure competitive growth rates (Bushell et al. 2017) in blood and identify genes involved in life-cycle progression (Ukegbu et al. 2023; Russell et al. 2023). However, the parasite in which genome-scale phenotypic screens were first applied was the African trypanosome, *Trypanosoma brucei (Baker et al. 2011; Schumann Burkard et al. 2011)*. In this organism, high-throughput approaches have been facilitated by a highly-effective system for inducible RNA interference (RNAi), which can be combined with methods for high efficiency transfection to screen mutants in a genuinely genome-wide fashion (Alsford et al. 2011).

RNAi-based screens in *T. brucei*, in particular a method known as RIT-seq (RNA interference target sequencing, (Alsford et al. 2011)), have a proven legacy of utility in situations where strong positive selection can be applied to select for RNAi effects, and have been used to identify genes associated with sensitivity to drugs (Baker et al. 2011; Schumann Burkard et al. 2011; Alsford et al. 2012; Baker et al. 2015; Collett et al. 2019), human serum components (Alsford et al. 2014; Lecordier et al. 2014; Currier et al. 2018), quorum sensing (Mony et al. 2014), and a wealth of cellular processes (Glover et al. 2016; Rico et al. 2017; Stortz et al. 2017; Trenaman et al. 2019; Marques et al. 2022). However, methods to date have substantial limitations in terms of quantitative power, how mutants are generated, and widening applicability to strains/species. One third of genes (951 of 2859) identified as having significant changes of fitness in RIT-Seq are associated with apparent increased growth rate (Alsford et al. 2011). Given that very few genes are expected to be associated with genuine gain of fitness on RNAi in culture, the vast majority of these genes will be false positives – implying that the true positive rate (sensitivity) in the pool of genes identified as significantly changed may be as low as 33% at Day 3, and lower still at Day 6 (25%; 1628 genes giving gain of fitness measurements). Such quantitative limitations are unimportant when selective screens can be used to isolate mutants with particular characteristics, but preclude its use as a quantitative tool for individual gene fitness.

Here, we present a ground-up redesign of parallel RNAi experiments in African trypanosomes, including generation of libraries optimized for NGS, direct amplification of fragments from parasites, and a new means of analysis. The result is a highly robust system for quantitative fitness cost measurement suitable for mapping fitness through individual components of cell complexes or processes. The system is transferable to different species of trypanosome and of sufficient sensitivity to show intra-species differences. We use this to quantitatively profile gene fitness in an strain of *T. brucei* widely used for infection studies.

## Results

### Direct-RNAi Fragment Sequencing

To develop a trypanosome RNAi library approach suitable for robust fitness quantification at genome-scale, we undertook a ground-up redesign of parallel RNAi experiments. We first created a library of RNAi fragments specifically optimized for next-generation sequencing by shearing whole genomic DNA from Lister 427 *T. brucei* and size selecting for fragments ∼400 bp. dsRNA of this size is sufficient to produce good RNAi in *T. brucei*, but is also suitable to be used directly for paired-end sequencing on Illumina platforms. This removes requirements for fragmentation, adapter addition and semi-specific PCR of parasite DNA as in RIT-Seq approaches, and means the identity of the entire RNAi fragment in each mutant can be inferred. Fragments were cleaned, tailed by linker addition and ligated into a version of a well-characterized RNAi construct p2T7 (LaCount et al. 2000) modified for high-complexity library generation (p2T7-V4δ, see Methods). The total complexity of the DNA library in bacteria is 7.6×10^6^ cfu (100x coverage of the genome). This method is extremely simple to apply to either whole genome DNA or sets of isolated of genes, making high-complexity strain, species, or gene-set specific libraries trivial to produce.

To generate tightly-regulated RNAi without variation due to differential silencing at individual rDNA loci, we selected a silent variant surface glycoprotein (VSG) locus at the subtelomere of a minichromosome (*VSG-31*) for integration of inducible RNAi constructs. This locus is stable with respect to mitosis and supports high-level transcription by T7 polymerase in a manner that can be regulated ∼400-fold (Wickstead et al. 2002). To use for genome-scale experiments, we created a redesigned construct to express an inducible, self-excising gene encoding I-SceI at this locus in a manner similar to that used at the rDNA locus (Glover and Horn 2009), but without the need to first tag an individual locus suitable for inducible expression. This construct was introduced into a trypanosome line stably expressing T7RNAP and TetR (SmOx-B4; (Poon et al. 2012)). Induction of I-SceI expression in these cells, and thus locus-specific double-strand break, robustly provides ∼30,000 independent transfectants in bloodstream-form Lister 427 under standard electroporation conditions in a clone-independent manner (Suppl. Fig. S1). Importantly, these I-SceI-inducible recipient cells are stable in the non-induced state to passage in culture and also freeze-thaw cycles without loss of function (Fig. S1), removing the need for the specific genetic background and fresh derivation of cells when using Sce* cells (Glover et al. 2015). We call these recipient cells 427-TTS (for *<u>T</u>*7RNAP, *<u>T</u>*etR, I-*<u>S</u>*ceI).

Two independent libraries each containing ∼300,000 independent RNAi mutants were generated in 427-TTS cells. Since the total size of the plasmid library is substantially greater (∼25-fold) than the number of trypanosome transfectants, the RNAi fragment present in a cell identifies both the genomic region targetted and also uniquely identifies the transfectant clone in the population in ∼99.9% of clones. To check this approach, we considered RNAi fragments that map to precisely the same genomic fragment but have the opposite orientation (and therefore must originate from different transfectants). Of 363,958 fragments across the 2 libraries mapping to the SMRT/Hi-C assembly of Lister 427 chromosomes (Müller et al. 2018), only 40 pairs have this relationship to other fragments in the same library – all mapping to DNA repeats in the genome (MAPQ ≤ 1) where the high-copy number increases the possibility of independent clones containing indistinguishable fragments. This confirms that sequencing of both ends of fragments in these libraries distinguishes individual RNAi clones in the population. Mapping fragments in this way allows the effect of RNAi in individual clones to be followed post-induction with full information regarding the RNAi fragment contained (Fig. 1B,C). In addition, it allows discrimination between fragments that uniquely map within the genome and those which might be derived from more than one place in the haploid assembly, including when different types of fragments are associated with the same gene (see Fig. 1C). It is worth noting that for experiments where the fragment library complexity is insufficient to uniquely identify clones, incorporation of additional barcodes during library linker addition is trivial.

**Figure 1.**
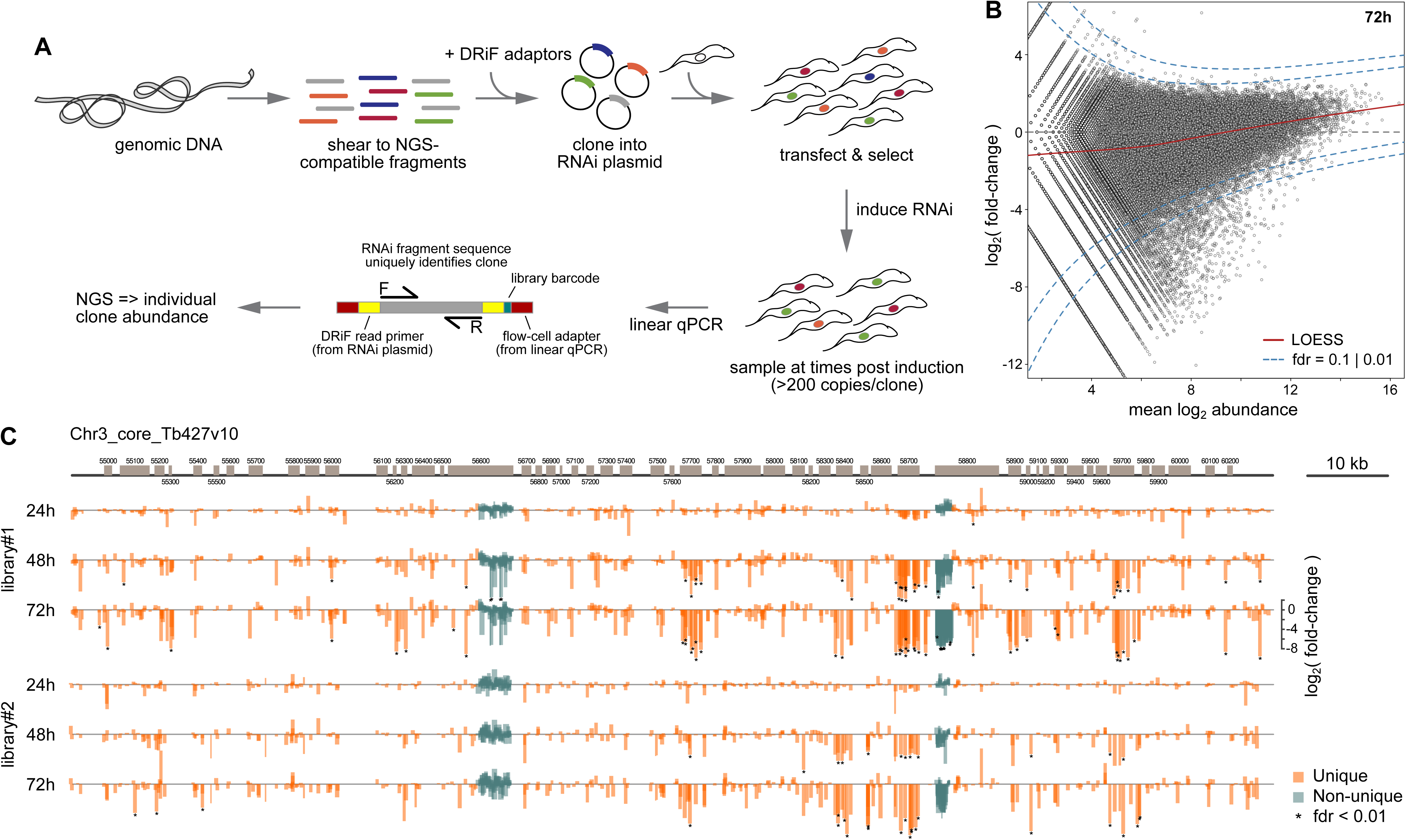
Direct-RNAi Fragment Sequencing for quantification of individual mutant effect sizes. A) Schematic of steps in production and sequencing of DRiF-Seq libraries. B) Example of analysis of 149,877 RNAi fragments in a single DRiF-Seq library based on fold-change and mean abundance of individual fragments. C) Fold-change for DRiF-Seq fragments mapping to a typical region of chromosomal ‘core’. Distinction is made between fragments which map uniquely to the assembly and those which could derive from more than one locus. Since the RNAi fragment is sequenced without fragmentation, the position and extent of each fragment is known, with uniquely mapping fragments representing individual transfectants followed through the course of each experiment. Individual fragments with fdr ≤ 0.01 are indicated.

In summary, significant modifications to previous RNAi-library approaches encompassed herein are: 1) a fragment library optimized for NGS; 2) integration at a stable minichromosomal locus with high transcription in the induced state, but very low transcription in the absence of induction; 3) an inducible recipient line (427-TTS) stable to freeze/thaw and growth in culture; 4) a new RNAi vector designed to take NGS-optimized fragments and incorporating custom read primers; 5) direct, quantitative amplification of fragments from parasite genomic DNA to avoid sub-sampling and saturation artefact. We call this combined method DRiF-Seq (*<u>D</u>*irect *<u>R</u>*NA*i <u>F</u>*ragment sequencing).

### Robust fitness change estimation by *a posteriori* noise estimation

A critical difficulty in analysis of induced mutant fitness in most RNAi libraries is that they do not conform well to conditions that allow robust estimation of dispersion (variance). Variation in such experiments arises from multiple sources, including: representation of mutants for a gene in the DNA library, the efficiency and timing of transfectant production, stochastic effects during library passage, differential effects of different mutants targeting a specific gene, the number of cell genomes sampled, and depth of sequencing. The latter is trivial to estimate, and in methods such as RNA sequencing without amplification, provides a good estimator of stochastic effects and gene-wise dispersion estimates which may be further moderated using weighted conditional likelihood (Robinson et al. 2010) or modelling (Love et al. 2014). However, for sequencing of amplicons (such as in Bar-Seq, RIT-Seq and CRISPR screens) read depth has little experimental meaning, and analysis tools designed for other applications perform poorly. In Bar-Seq for gene knockout in *Plasmodium*, a careful estimation of variation introduced by each step was used, with replication across transfection pools to provide estimates of error associated with mutant abundance at each point of the experiment that were subsequently applied to calculation of relative growth rates (Bushell et al. 2017). However, such approaches require a lot of information about individual steps that is often difficult to estimate. Moreover, for a library of randomly-sheared fragments, the same RNAi mutant is highly unlikely to occur in independent libraries, and even large experimental set-ups only allow small numbers of full replicates (usually 2-4), making variance of individual genes a poor estimator of true dispersion.

As an alternative approach, we used the distribution of DRiF-Seq data in the experiment itself to model the statistical parameters. These data contain stochastic contributions produced by measurement noise or other random sources plus true signal. We used a simple feature of DRiF-Seq data – that the data will include a substantial proportion of points (e.g. non-genic RNAi fragments or fragments targetting non-essential genes) – to make *a posteriori* estimates of signal (effect size) and noise (dispersion) for all data points. The only assumptions made in this approach are that: 1) all sources of noise will sum to an approximately Gaussian distribution for a given count depth, and 2) very few knockdowns are likely to increase cell fitness. Note that this latter assumption does not exclude the possibility of knockdowns that cause faster growth, but uses the fact they will be in the great minority.

One advantage of *a posteriori* noise estimation (*ap*NE) is that it can estimate the significance of changes in abundance for RNAi mutants which occur only once across libraries (as nearly all mutants do) without the need to assume their effect is the same as other mutants in the same gene. Figure 2A-C demonstrates the application of *ap*NE to 214,083 mappable RNAi fragments from a single DRiF-Seq library in Lister 427. The method can also be applied to gene-wise abundance counts (Fig. 2D) providing excellent quantitation of effect sizes and statistical discrimination between likely genuine changes and noise. Gene significance correlation with DRiF-Seq and *ap*NE was very good between entirely independent mutant libraries (Fig. 2E; effect size ρ=0.74), as is the measured effect of the average independent mutant to the measurement made by combining the abundance of all (Fig. 2F; effect size ρ=0.90). Raw counts and *ap*NE estimates of fitness cost and confidence for 363,958 mappable fragments are provided in Suppl. Data Files 1 and 2, and gene-wise estimates are provided in Suppl. Data File 3.

**Figure 2.**
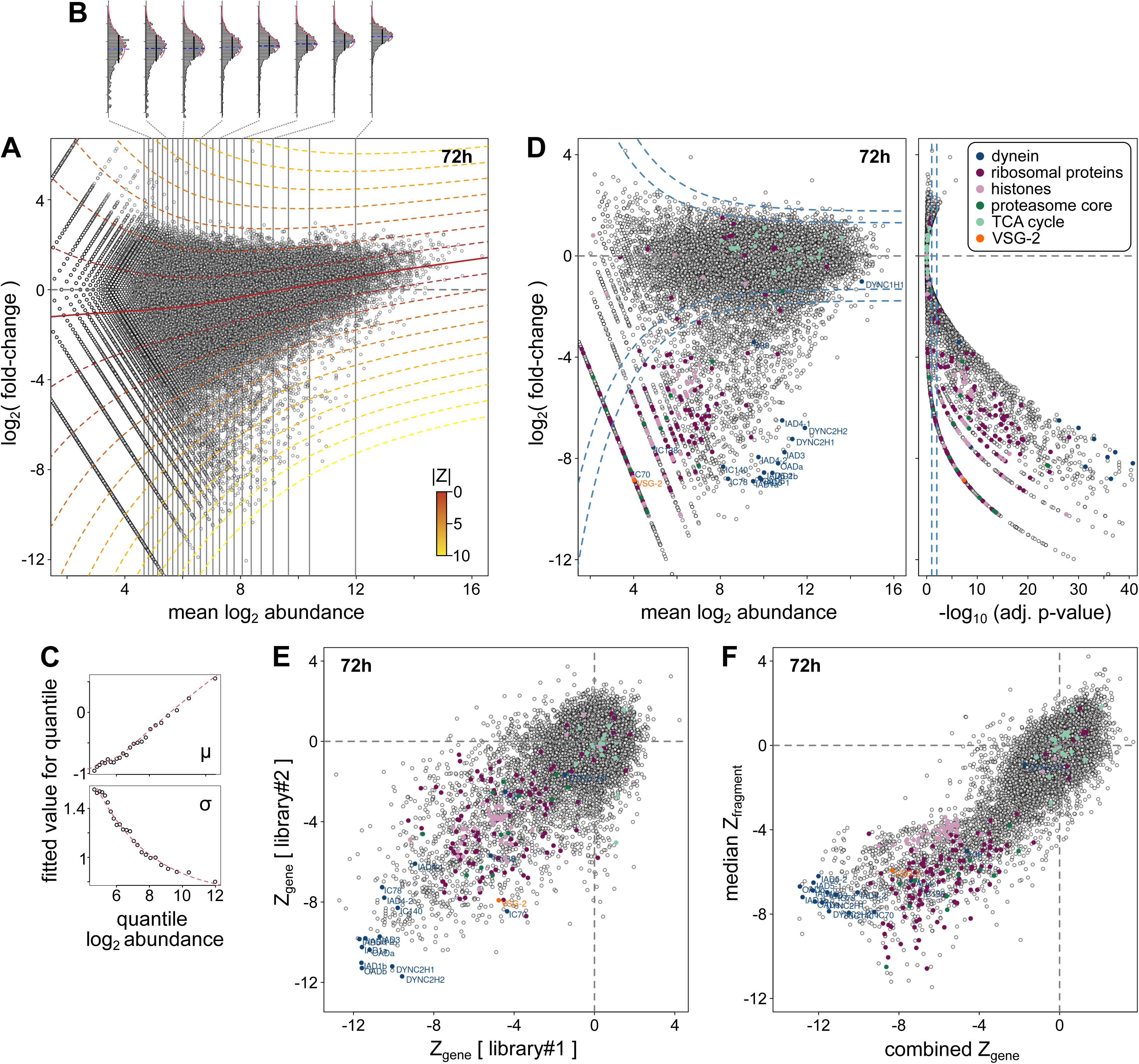
Analysis of individual fragment and gene-wise effect sizes by *a posteriori* noise estimation. A) MA-plot showing changes in abundance for 149876 mapping fragments (count ≥ 24) in a single DRiF-Seq library in Lister 427 parasites. Since most fragments will not be associated with gain of fitness, a good estimate of dispersion at a given abundance can be estimated by fitting the top 75% of data at each abundance quantile to a censured Gaussian (B). The distribution of these parameters across abundance quantiles (C) allows the calculation of Z scores for individual fragments based on their abundance change and the normal dispersion of data at this abundance. D) MA- and volcano plot showing gene-wise changes in abundance for 95187 fragments mapping to the CDS of 14758 (of 17243) annotated genes. Lines representing fdr = 0.1 and 0.01 are shown. There is a very high correspondence between calculated Z-scores across libraries (E) and between Z-scores calculated for the total gene count or the median fragment Z (F).

These data demonstrate that DRiF-Seq can be used to generate reproducible and quantifiable RNAi mutant effect size estimates at both gene- and individual mutant-level. The associated *ap*NE implementation may also be applied to previously generated data, robustly separating likely artefactual calls of significance in RIT-Seq data introduced by DESeq (Suppl. Fig. S2). It is also widely applicable to data from other genome-scale functional screens based on sequencing (e.g. Bar-Seq, siRNA libraries, CRISPR screening, etc.) – for example, reanalysis of data from a genome-wide CRISPR screen in *Toxoplasma (Sidik et al. 2016)* readily provides statistical treatment for the average effect of sgRNAs targetting each of 8158 genes, gives an estimate for the enrichment of genes with a null phenotype in the experiment (1.42-fold), and demonstrates that ∼47% of genes have detectable loss-of-fitness effects in vitro upon targetting by Cas9 (Suppl. Fig. S3). It also identifies the 2800 genes associated with a significant loss-of-fitness (fdr ≤ 0.01) in this screen, shows that there are no gain-of-fitness genes at the same threshold, and in addition provides false-discovery rate estimates for each of the 81,389 individual sgRNAs detected. *ap*NE re-analysis of RIT-Seq and CRISPR screen data are available in Suppl. Data Files 4 and 5, respectively.

### Genome-scale DRiF-Seq in quorum-sensing competent trypanosomes

One advantage we anticipated for the DRiF-Seq approach was the ability to transfer into different genetic backgrounds. We applied the DRiF-Seq method to EATRO1125 strain trypanosomes (expressing VSG AnTat1.1E) – a pleomorphic cell line that is mammal and fly transmissible (characteristics that have been lost in Lister 427), but has not been amenable to RNAi library approaches due to lack of proven genetic tools and low transfection efficiency. Creation of library-compatible “AnTat-TTS” cells required only two transfections – introduction of pSmOx (Poon et al. 2012) and then recreation of the inducibly cleavable *VSG* target site on a suitable minichromosomal locus (allowing the same library plasmid to be introduced into either strain). The modified line behaves like wild-type AnTat1.1E cells in culture (Fig. 3A), exits the cell cycle in response to quorum sensing (Fig. 3B), and makes characteristic waves of parasitaemia in animal models with no significant change in virulence (Fig. 3C,D), making it a suitable chassis for studying important aspects of trypanosome infection biology both in vitro and in vivo. As for L427-TTS cells, AnTat-TTS is stable to freeze-thaw and growth in culture, and induction of I-SceI expression allows integration of p2T7-V4δ at ∼4000 independent transfectants per standard transfection (Suppl. Fig. S1D).

**Figure 3.**
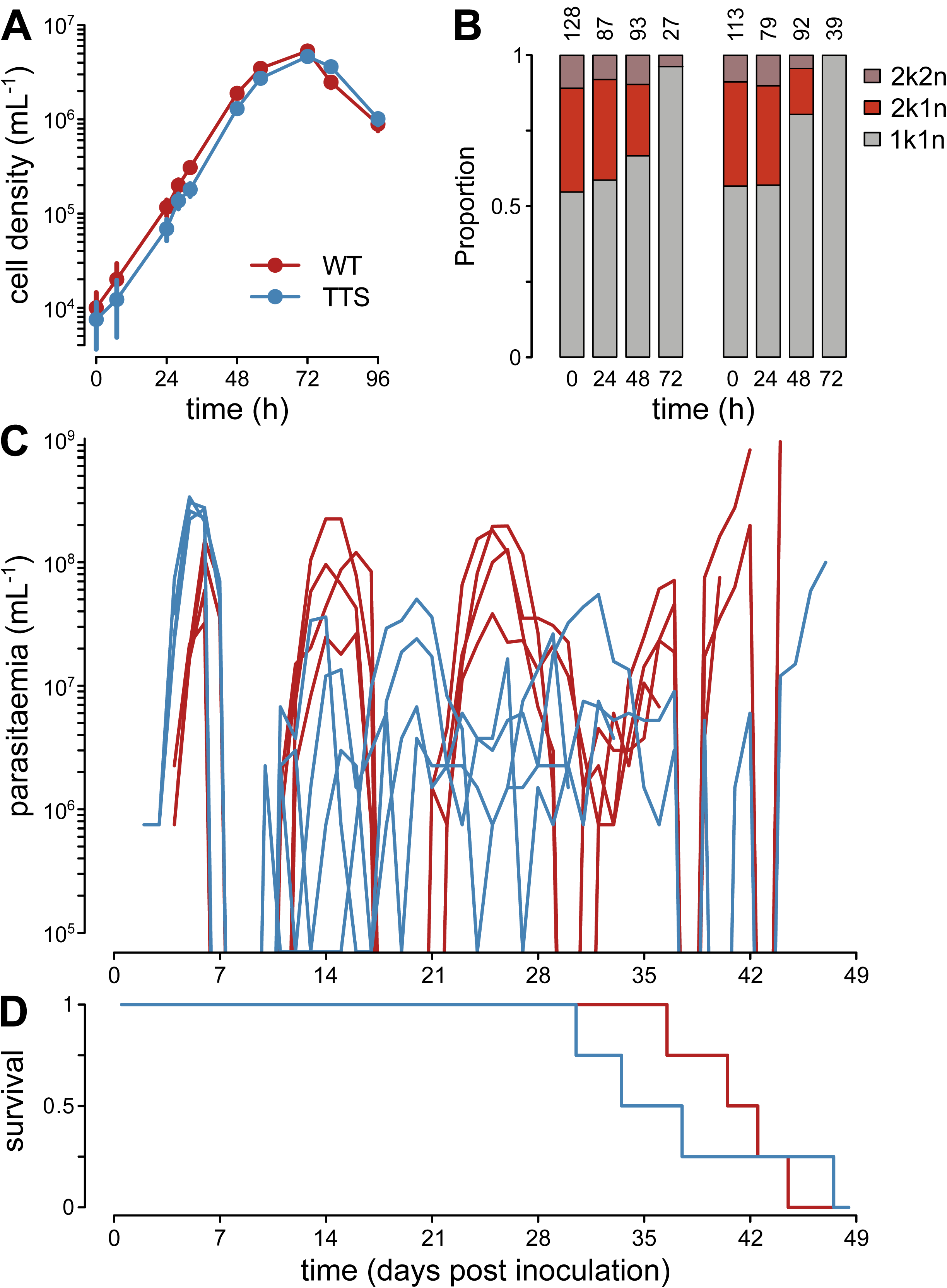
Development of a DRiF-Seq-compatible line in pleomorphic trypanosomes. AnTat-TTS (based on bloodstream-form EATRO1125 cells) expresses TetR and T7RNP for inducible expression and a stable self-excising cassette at a minichromosomal locus which can be targetted at high-efficiency with DRiF-Seq constructs used for Lister 427. Cells following genetic modification grow indistinguishably from unmodified EATRO1125 cells (A) and exit the cell cycle in response to cell density (B). In infection models, AnTat-TTS recreate parasitaemic waves seen for unmodified pleomorphic cells (C) and have no significant change in virulence (D).

We created a strain-specific fragment library using EATRO1125 genomic DNA in the p2T7-V4δ vector and used this to generate a population of ∼260,000 independent RNAi mutants in AnTat-TTS. To investigate fitness of gene knockdowns during growth in culture, RNAi was induced in this population whilst keeping cell numbers below the density necessary to stimulate quorum sensing at all times. To aid comparison with DRiF-Seq experiments in Lister 427, RNAi fragments were mapped to the SMRT/Hi-C assembly of Lister 427 chromosomes (Müller et al. 2018). However, the total number of mappable fragments above threshold increases only marginally if the recent long-read assembly of EATRO1125 is used as reference ((Naguleswaran et al. 2021); 260,621 and 263,694 mappable fragments using Lister 427 and EATRO1125 assemblies, respectively) indicating that little usable information is being lost by this cross-mapping. Raw counts and *ap*NE analysis of mappable fragments and genes are provided in Suppl. Data Files 6 and 7.

In spite of a longer doubling time in culture (7.5 h and 6.5 h for EATRO1125 and Lister 427, respectively), growth defects upon RNAi knockdown are greater in EATRO1125 and have a substantially lower variance (Fig. 4B; mean 10-fold difference in induced/non-induced ratio at 72h post-induction for genes with a loss-of-fitness in both). Whether this reflects more active RNAi machinery or some other biological factor is currently unknown, but it means that DRiF-Seq in EATRO1125 provides further quantitative improvement for gene mutant fitness over that already seen in Lister 427. >5000 genes cause a significant detectable loss-of-fitness when targetted by RNAi in EATRO1125 at a 10% false discovery rate. As *ap*NE provides an explicit estimation of experimental noise, it is possible to also estimate over-/under-representation of genes outside of the expected distribution (Fig. 4C) even when confidence of specific gene contributions would be below a suitable threshold. This demonstrates that >50% of all ‘core’ genes (non-VSG genes occurring in the diploid regions of the assemblies) are associated with a loss-of-fitness when knocked down in bloodstream-form *T. brucei*, of which 3601 (44% of 8170 targetted) can be identified with very high confidence (fdr ≤ 0.01). This suggests that even in a life-cycle stage where important aspects of cell biology (such as the mitochondrial function) are heavily down-regulated due to scavenging of material from the host, African trypanosomes still rely on a large proportion of their genetic repertoire. Moreover, the proportion of genes with fitness costs when targetted by RNAi is very similar to the *ap*NE estimate for *Toxoplasma* based on Cas9 targetting, but with a much greater range of effect sizes being quantifiable (Fig. 4D). In comparison to loss of fitness, very few genes are associated with an increase in cell number (gain-of-fitness) on RNAi in DRiF-Seq (for example, only 12 core genes have apparent gain-of-fitness at 6 days post-induction with fdr ≤ 0.01; Fig. 4C). This is in contrast to the high proportion of apparent gain-of-fitness phenotypes seen in RIT-Seq profiling (Fig. 4E) and agrees with the strong expectation that there are likely very few genes actively suppressing growth at densities below those necessary to stimulate quorum-dependent growth arrest (‘stumpy’ formation).

**Figure 4.**
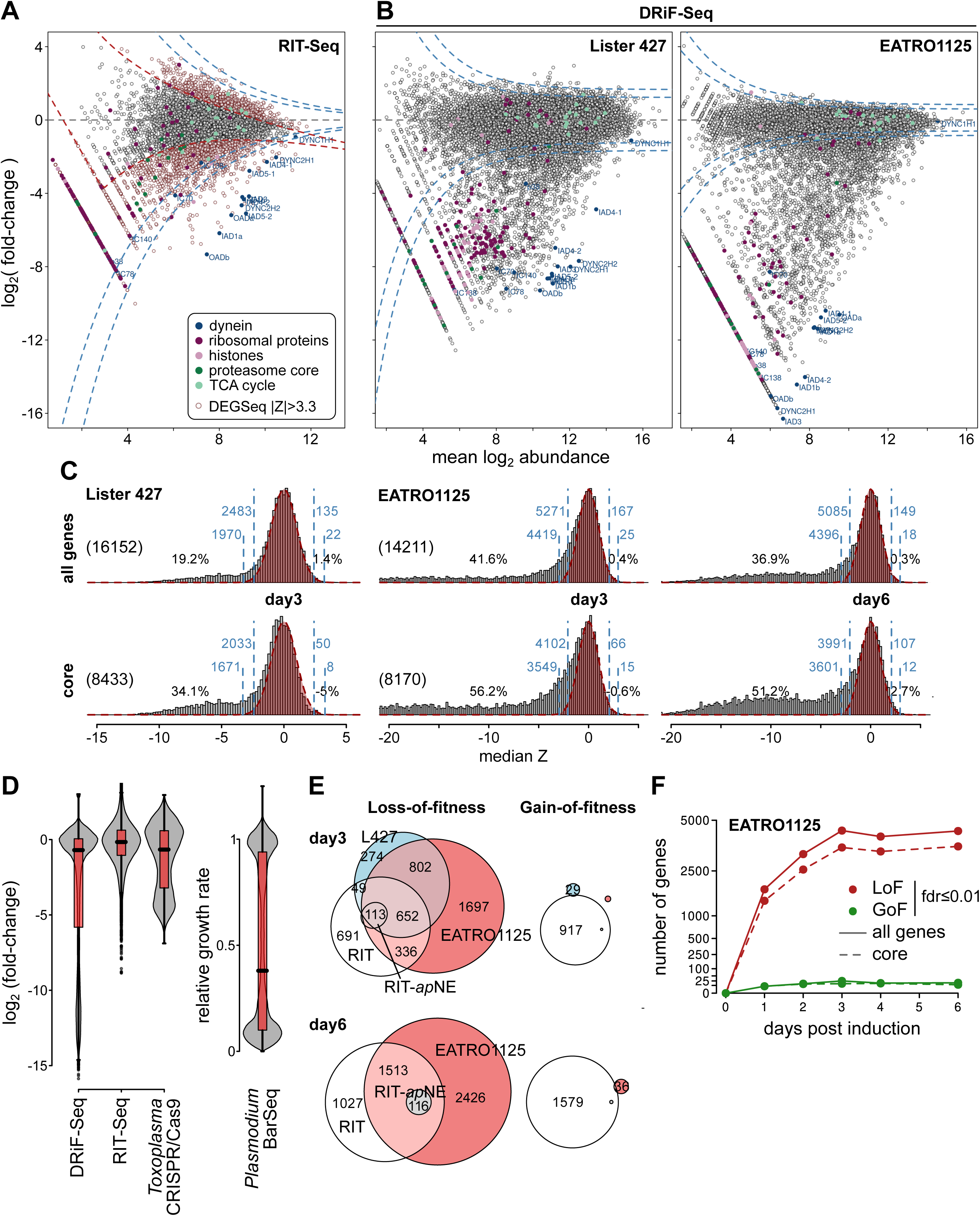
DRiF-Seq provides robust fitness change estimates for genes in pleomorphic trypanosomes. A) MA-plot for data from previous high-throughput phenotyping of genes in Lister 427 (RIT-Seq; 7435 non-redundant TREU927 genes (Alsford et al. 2011)). Genes identified as associated with a significant gain- or loss-of-fitness in previous analysis by DEGSeq (|Z| > 3.3) are indicated (red) alongside reanalysis of the data by *a posteriori* noise estimation (blue lines at fdr = 0.1 and 0.01). B) MA-plots for data from DRiF-Seq libraries in Lister 427 (16152 genes) or EATRO1125 (14211 genes) showing *ap*NE as in (A). Data in (A) and (B) are for day 3 post-induction. Genes encoding dynein heavy, intermediate or light chains, components of the ribosome, histones, proteasome core, and enzymes of the tricarboxylic acid cycle are highlighted. C) Distribution of median fragment Z scores for DRiF-Seq libraries in Lister 427 and EATRO1125 at days 3 and 6 post-induction. Data are shown for all genes or only non-*VSG* genes in the diploid ‘core’ genome. The expected null distribution estimated from the fitted variance of the data is overlaid in red. The number of genes detected by *ap*NE as enriched/depleted at fdr = 0.1 or 0.01 are shown in blue, in addition to percentage over-representation of genes in each distribution tail. Total number of genes analysed is given in parentheses. D) Comparison of genome-wide gene-ablation effect sizes in DRiF-Seq and RIT-Seq in *Trypanosoma* with those from CRISPR/Cas9 disruption in *Toxoplasma (Sidik et al. 2016)*, and relative growth rate of 2578 full gene knockouts in *Plasmodium (Bushell et al. 2017)*. E) Number of genes detected as enriched (GoF) or depleted (LoF) by DRiF-Seq at fdr ≤ 0.01 over 6 days of induction. F) Comparison of gene-sets classified as having significant loss- or gain-of-function in DRiF-Seq in Lister 427 (L427) and EATRO1125 cells (using *ap*NE) with original calls made in Lister 427 by RIT-Seq (RIT; DEGSeq |Z| > 3.3). Calls following reanalysis of the RIT-Seq data with *ap*NE are also shown (RIT-*ap*NE). For clarity, numbers are shown for sets of >20 genes only.

Notably, the DRiF-Seq method is able to detect most deviation from normal growth very quickly after induction – for example, 1427 core genes can be assigned a highly significant LoF even at 24h post-induction, and the number of genes associated with significant changes in fitness on RNAi at reliable thresholds (fdr ≤ 0.01) does not increase after day 3 of induction (Fig. 4F). This does not represent reversion of RNAi effects, which continue to accumulate over time (see below). However, it does demonstrate that most RNAi defects (at least for bloodstream-form EATRO1125) occur quite rapidly post-induction and there are limited benefits to longer induction times for most genes, as greater effect sizes are offset by increased stochastic noise present in samples at higher passage numbers.

### Mapping fitness landscapes within molecular complexes

Quantitative enrichment analysis shows that most gene sets annotated with gene ontology (GO) terms have mean negative effects on growth when targetted in DRiF-Seq (Fig. 5A, Suppl. Fig. S4), as expected for functionally annotated genes. Sets associated with core cell functions such as translation, flagellar function, vesicle movement, nuclear import/export and cell division have significantly higher LoF effects than average effects for genes with GO annotations (Fig. 5A). Comparison of LoF effects across different subcellular locations from systematic localisation data (Billington et al. 2023) also agrees well with known trypanosome biology, with low median effects for genes targetted to the mitochondrion or plasma/flagellar membrane in insect-form cells, but large LoF for most gene products localised to the nuclear pore, kinetochore or intraflagellar transport particles (Suppl. Fig. S5). Similar trends in GO term effects are seen in data from RIT-Seq experiments (Alsford et al. 2011; Horn 2022), but the quantitative data here are in very good agreement with known trypanosome biology – including lack of LoF when targeting many aspects of mitochondrial function (including the TCA cycle), β-oxidation and de novo pyrimindine biosynthesis (which is not essential when extracellular pyrimidine is available (Ali et al. 2013)). However, the quantitative capacity of DRiF-Seq provides an opportunity not only to look at mean effect sizes for GO terms *en masse*, but also to map fitness landscapes for individual components within defined complexes/structures (Fig. 5B). There is a remarkable consistency of measured fitness cost when targetting components of many complexes. For example, targetting the 2 largest subunits that define the different RNA polymerase complexes gives very similar effects, the magnitude of which is characteristic for each polymerase class (Fig. 5B). Similarly, components of the different clathrin-associated adaptor protein complexes have a clear hierarchy of fitness cost (AP-1 > AP-3 > AP-4), with ablation of AP-4 having no detectable effect on growth. In agreement with the inferrence that AP-4 likely has a limited role in *T. brucei*, all subunits of this complex are absent in the genome of the related African trypanosome, *T. congolense*. As anticipated for bloodstream-form trypanosomes where mitochondrial metabolism is highly reduced, many proteins associated with the mitochondrial inner membrane also have no loss of fitness when targetted in EATRO1125. This includes Complexes I and II of the electron transport chain (Fig. 5B), matching inference from the viability of knockouts of 2 components of Complex I (Surve et al. 2012) and RNAi targeting SDH1 (Alkhaldi et al. 2016), but here extended across the entire complexes. In infective EATRO1125 cells, one of 36 subunits specific to Complex I and II targetted in 160 independent mutants gave a statistically detectable LoF across 18 cell generations, whereas other inner membrane proteins such as components of the TIM translocase complex have a clear significant LoF.

**Figure 5.**
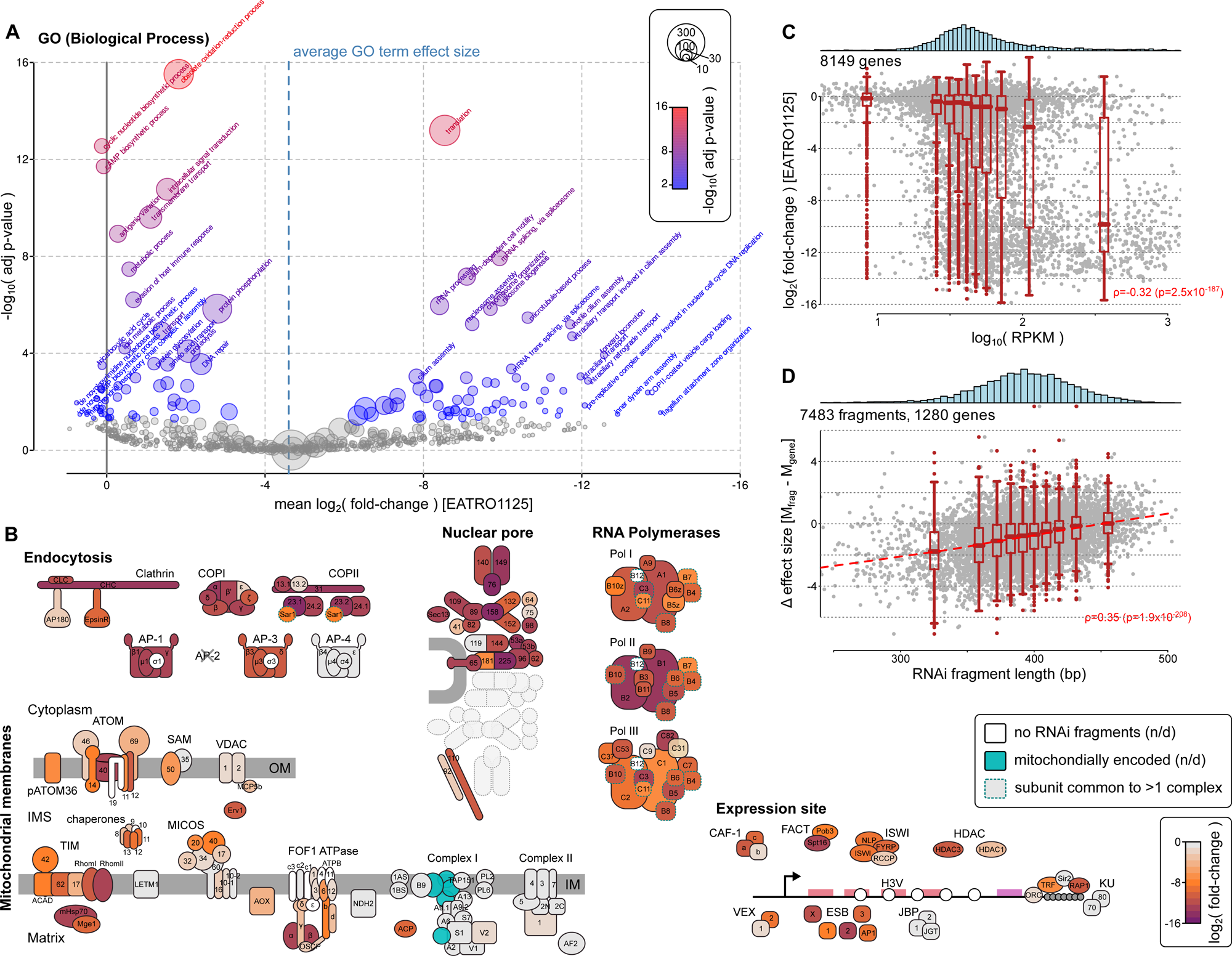
DRiF-Seq allows quantitative mapping of fitness landscapes. A) Quantitative enrichment analysis of GO Terms highlighting Biological Process terms that have fitness effect sizes significantly different (adjusted p-value ≤ 0.05; Monte Carlo permutation test) from the average effect size of genes with GO annotation. Circle size indicates number of measured genes associated with each GO term. Data for Cellular Component and Molecular Function ontologies are shown in Suppl. Fig. S4 and full data are available in Suppl. Data File 8. B) Fitness landscapes across proteins involved in five key cellular processes/locations in trypanosomes. Components which had no mapping RNAi fragments in DRiF-Seq libraries are not filled. Components which are encoded by mitochondrial DNA and unable to be targetted by RNAi are also highlighted (light blue). The AP-2 complex is not present in African trypanosomes *(Manna et al. 2013)*. Subunits common to more than one polymerase are indicated by dashed outline. C) DRiF-Seq effect size versus gene transcript levels (RNA-Seq reads per kilobase per million reads, RPKM). Reads for bloodstream-form EATRO1125 cells were taken from European Nucleotide Archive study ERP125374. D) Length-dependent differential effect sizes for fragments targetting 1280 genes associated with large loss-of-fitness (≥ 256-fold depletion; fdr ≤ 0.01) and targetted by ≥ 2 fragments. Difference between calculated effect size of individual fragment versus overall gene loss-of-fitness is shown. All data in (A-D) are for effect sizes at day 6 post-induction in EATRO1125.

Quantitative fitness data can also rapidly identify likely dispensible components of essential complexes/processes (Fig. 5B) – for example, Nup119 and Nup75 are not associated with significant LoF, despite the central location in the nuclear pore of the former (Obado et al. 2016), and targetting the Pol III subunit RPC9/RPC11 has no significant loss of fitness, mirroring the low cost of ablation of the vertebrate homologue (Wei et al. 2016). These data also distinguish between likely essential and non-essential paralogue pairs (e.g. the 2 paralogues of SEC13 predicted to be part of the COPII complex in trypanosomes). To our knowledge, none of these effects have been demonstrated in a systematic manner before this work.

There is a very highly significant correlation between LoF effect size and mRNA levels in bloodstream form cells (Fig. 5C), but this reflects only that many highly-expressed genes are associated with essential cell functions rather than expression level being a good predictor of fitness cost on knockdown or necessary for RNAi effect – there are both very highly expressed genes which still have no LoF on knockdown and also genes with very low mRNA levels associated with substantial fitness costs. As DRiF-Seq maps fitness of individual mutants for which we know the precise RNAi fragment, these data also contain information about the effectiveness of RNAi in *T. brucei*. RNAi targetting CDS have significantly higher loss-of-fitness than fragments targetting 3’-UTR of the same genes (Suppl. Fig. S6A), but this may reflect differences in the confidence of UTR versus CDS annotation or effects of alternative splicing (where shorter 3’UTRs might not be targetted). It is unlikely to reflect the effects of different composition, as although fragments with higher GC do tend to have a greater effect size than lower GC fragments targetting the same gene, the effect is very modest (Suppl. Fig. S6B). Surprisingly, however, the data show convincingly that RNAi fragments at the smaller end of the length distribution in our DRiF-Seq libraries (∼300 bp) give a greater LoF than larger fragments targetting the same gene (Fig. 5D). The effect equates to an average of ∼1.5 fewer cell generations across 6 days of induction for genes with significant LoF, and is extremely robust (p = 10^-79^), but has important implications for the design of RNAi experiments in trypanosomes. Unsurprisingly, mRNA stability may also affect RNAi effect (Suppl. Fig. S6C), but the size of this effect cannot be deconvolved from the fact that there are more essential genes and fewer hypotheticals amongst the transcripts with longer half-lives (Suppl. Fig. S6D).

### DRiF-Seq reveals biological differences at subspecies resolution

Our data show that the DRiF-Seq method is both sensitive and robust for estimation of quantitative fitness costs of individual gene knockdowns in *T. brucei*. Lister 427 and EATRO1125 cells differ in some key aspects of transmission biology, but are also very closely related strains of the same parasite sub-species. We reasoned that, if DRiF-Seq were sufficiently sensitive to predict differences in biology between even such closely-related strains, then quantitative fitness estimates from DRiF-Seq are highly likely to be useful in many other experimental contexts.

To test this, we compared estimated knockdown effect sizes in Lister 427 and EATRO1125 at 72 h post-induction of RNAi (Fig. 6A). To avoid known differences in quorum-sensing between the strains, both were grown below the density inducing ‘stumpy’ formation at all times. As already seen, the effect size for knockdown of most essential genes is greater in EATRO1125 than Lister 427 (Fig. 6A). However, there are substantial numbers of genes that have an apparent differential effect when targetting in one strain even above the general trend, including genes which are essential in Lister 427 but have no measured effect when targetted in EATRO1125 in spite of the greater average RNAi effect (e.g. procyclin-associated genes PAG2 and PAG5; Fig. 6A). These differences do not appear to be the consequence of targetting different parts of a gene in the 2 strains, as the differential effects of greater loss-of-fitness in either EATRO1125 or Lister 427 replicated in both direction and magnitude in all cases when targetted with the same fragment in independently-derived knockdown lines (4 and 6 genes, respectively; Fig. 6B and Suppl. Fig. S7). This suggests that most differences in knockdown observed between the strains represent genes that are functionally more sensitive to knockdown in one strain relative to the other.

**Figure 6.**
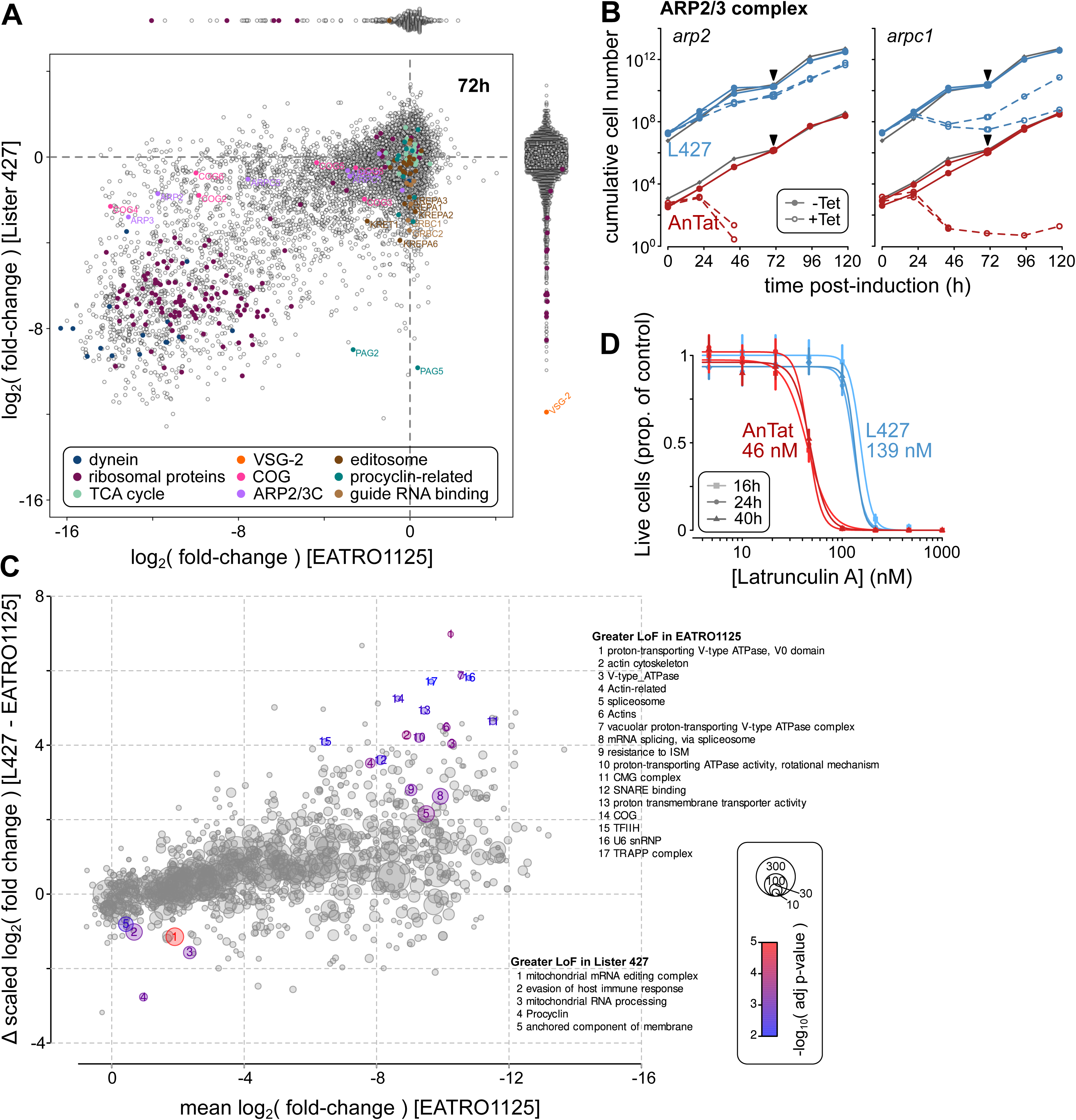
Differences in sensitivity to depletion of biological processes between two strains of *T. brucei*. A) Comparison of DRiF-Seq effect sizes for genes when targetted in Lister 427 or EATRO1125. For comparison, reads from both strains were mapped to the Lister 427 SMRT/Hi-C assembly (Müller et al. 2018). Effect sizes for genes targetted in one strain but not the other are shown in the beeswarm plots above and to the right of the main plot. B) Comparison of growth of Lister 427 (L427) and EATRO1125 (AnTat) cells in the presence (+Tet) or absence (-Tet) of induction of RNAi targetting components of the ARP2/3 complex. For comparison, cumulative cell numbers of the 2 strains have been offset by 10^2^. Grey lines show growth of parental lines not carrying RNAi construct. C) Differential mean effect size between Lister 427 and EATRO1125 for 1588 GO terms and gene sets associated with 8283 genes. Data are scaled to account for the overall greater effect size of knockdown seen in EATRO1125. Terms behaving significantly differently than bulk are highlighted (adjusted p-value ≤ 0.01; Monte Carlo permutation test). Circle size indicates number of measured genes associated with each GO term. Full data are available in Suppl. Data File 9. D) Comparison of sensitivity of wild-type Lister 427 (L427) and EATRO1125 (AnTat) strains to treatment with Latrunculin A for 16, 24 and 40h. Curves show least-squares fit of each dataset to Hill equation. Inferred EC_50_ concentrations for each line are also given next to curves. Bars: s.e.m.

Quantitative enrichment analysis of differential knockdown effects for GO terms and gene-sets (scaled for the average behaviour of genes in each strain) shows there is a selective bias in the genes showing these differential sensitivity in Lister 427 and EATRO1125 cells (Fig. 6C). A total of 17 and 5 terms are highly significantly enriched with respect to greater loss-of-fitness effect in EATRO1125 and Lister 427, respectively. Production of proteins in the mitochondrion is more sensitive to knockdown in the more culture-adapted Lister 427 line. In contrast, EATRO1125 is significantly more sensitive to knockdown of genes associated with the actin-based cytoskeleton and aspects of ER/Golgi function and splicing (Fig. 6A,C). This line also has greater sensitivity to perturbation of V-type ATPase function (Fig. 6C), which has important consequences for generation of resistance to the important veterinary drug isometamidium (Baker et al. 2015).

Given that we observe DRiF-Seq effect biases across multiple genes, they are highly unlikely to represent simple differences in RNAi, but genuine differences in the sensitivity of the strains to perturbation of the systems targetting. To test this, we perturbed one predicted differential system – the actin cytoskeleton – with available inhibitors (acting downstream of mRNA). In agreement with DRiF-Seq prediction, EATRO1125 cells are ∼3x more sensitive to latrunculin A inhibition of actin polymerization than Lister 427 (Fig. 6D). These data show that the robust fitness measurements available in our method can predict biological differences, including differential pharmacological sensitivity, even at individual strain level, suggesting it can in future be used to probe for biological differences between related species – and, importantly, between different conditions, such as in infections versus culture.

### Comparative functional genomics across eukaryotic lineages

As we were able to use DRiF-Seq fitness profiles to find biological differences at a sub-species level, we next tested whether this could be applied at much larger evolutionary scales. Genes more widely conserved across eukaryotes are predicted to be more often associated with essential processes. In agreement with this, more widely-conserved genes tend to have higher fitness costs when targetted in DRiF-Seq libraries (Fig. 7A). However, comparative genomics alone is not a substitute for quantitative loss-of-fitness (at least in terms of orthologue distribution) and many genes that are found only in African trypanosomes have strong effect sizes when knocked down, while genes common to eukaryotes are not necessarily essential (Fig. 7A) – demonstrating the need for functional comparisons in addition to genomic ones.

**Figure 7.**
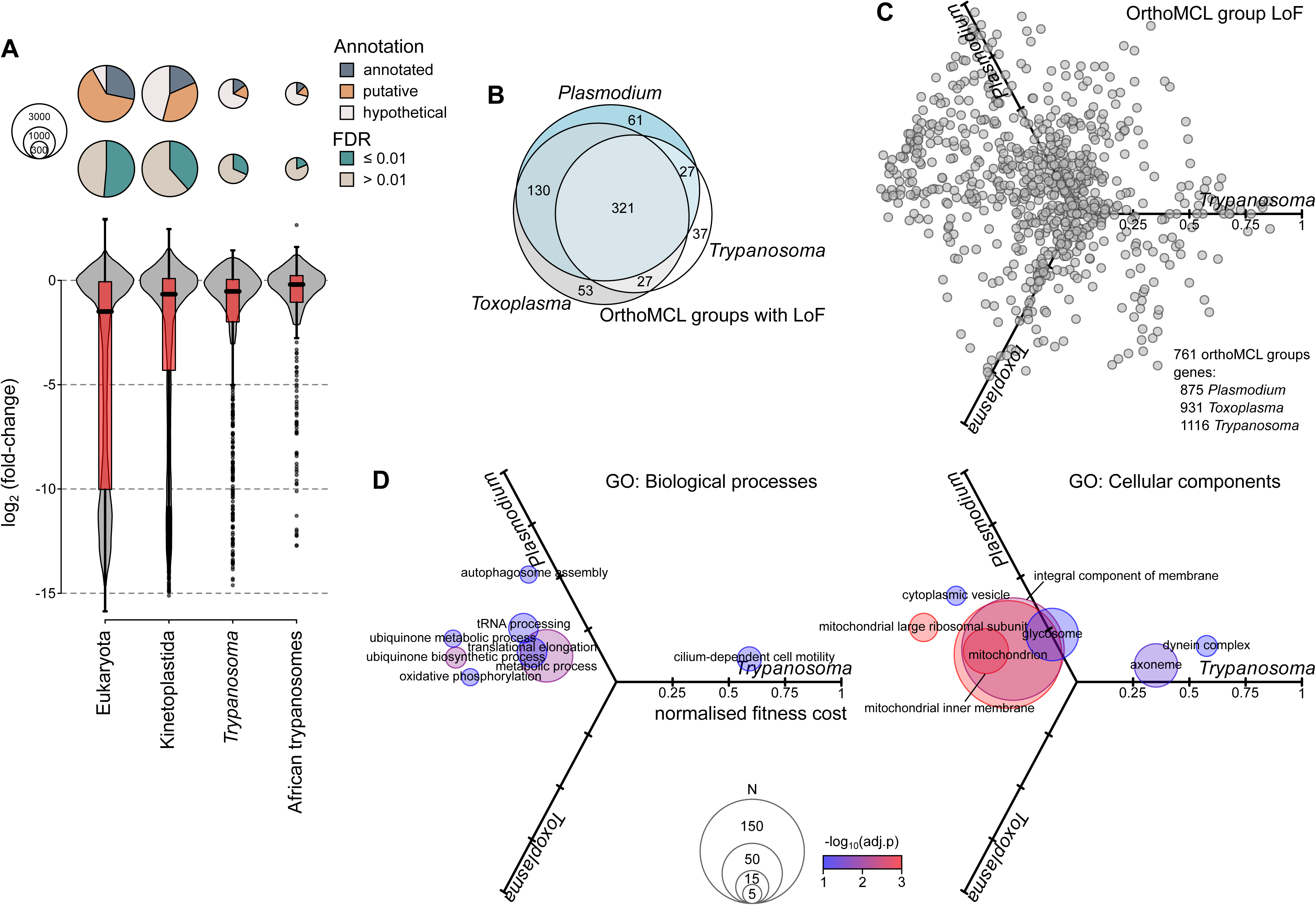
Comparative functional genomics across eukaryotic lineages. A) Effect sizes for trypanosome genes belong to orthoMCL groups found across non-kinetoplastid eukaryotes (Eukaryota) or restricted to the class Kinetoplastida, genus *Trypanosoma* or only the African (salivarian) trypanosomes. The proportion of genes in each class with functional annotation, putative function, or annotated as ‘hypothetical’ (no assigned function) are shows above along with proportions associated with significant change of fitness (fdr ≤ 0.01). B) Intersection of fitness costs for genes from 656 orthoMCL groups having a detected loss-of-fitness (normalised mean fitness cost of ≥ 0.3) in at least one organism. C) Radial plot of normalised loss-of-fitness effect size for 761 orthoMCL groups common to *Trypanosoma*, *Toxoplasma* and *Plasmodium* and including genes for which quantitative fitness data have been measured in all 3 organisms. Position is mean effect size for all genes associated with orthoMCL group in each organism. D) Mean positions of GO terms associated with 761 shared orthoMCL groups on radial axis from (C). Only GO terms which are significantly displaced distally (adjusted p-value ≤ 0.1; Monte Carlo permutation test) are shown. Circle size reflects the total number of genes encompassed by each GO term.

As large-scale quantitative fitness data exist for *Plasmodium* (knockouts for 2578 genes; (Bushell et al. 2017)), *Toxoplasma* (Cas9-targetting of 8158 genes; (Sidik et al. 2016)) and *Trypanosoma* (RNAi-mediated knockdown of 14,211 genes in EATRO1125 here), we compared the fitness costs for mutants in sets of orthologues existing between the 3 organisms. A total of 761 orthoMCL groups have genes with fitness profiles in all 3 organisms (encompassing 875, 931 and 1116 genes for *Plasmodium*, *Toxoplasma* and *Trypanosoma*, respectively), of which 656 have a loss-of-fitness detected in 1 or more organism (Fig. 7B). Half of these (49%; 321 of 656) have a LoF in all 3 organisms, in agreement with the general link expected between conservation and essentiality, with a further 20% (130) having a detected LoF in the 2 more closely-related apicomplexan parasites only.

Bias in fitness cost conforms well to known differences in biology between the parasites (Fig. 7C,D), including a dependence on axonemal motility for bloodstream-form trypanosomes (both *Plasmodium* and *Toxoplasma* cells having no flagellum in the stages tested) and a converse lack of sensitivity to perturbation of mitochondrial processes including oxidative phosphorylation and ubiquinone metabolism. In contrast, fitness costs from removal of genes associated with autophagy are significantly higher in *Plasmodium* than the other parasites (Fig. 7D).

### Kinetics of fitness loss map to biological functions

One great advantage of the system for RNAi in *T. brucei* is that selection of stable transformants is fully separated from induction of RNAi. This allows the timings of fitness costs to be investigated in addition to the fold-change at a specific end-point. To analyse a conservative set of genes associated with a likely fitness cost, we took 3806 genes associated with a significant loss-of-fitness at a stringent false discovery rate (fdr ≤ 0.01) from both total gene-count and median fragment effect. Principal component analysis of change in RNAi target abundance across 6 days of RNAi induction shows clear structure in temporal mutant fitness (Fig. 8A). Unsupervised clustering of genes based on their profile of log fold-change through the RNAi time series (Fig. 8B) partitions the data into 6 sets: 5 sets of genes with profiles consistent with ‘early’ to ‘late’ loss-of-fitness effects (Fig. 8C,D), and a 6^th^ set associated with approximately constant and relatively low loss-of-fitness (‘constant LoF’ in Fig. 8D). 11 genes associated with significant gain-of-fitness under the same criteria show an approximately constant increase in growth rate equivalent to reduction in generation time of ∼0.5-1 hour (Fig. 8D), again suggesting that these are genuine gain-of-fitness knockdowns.

**Figure 8.**
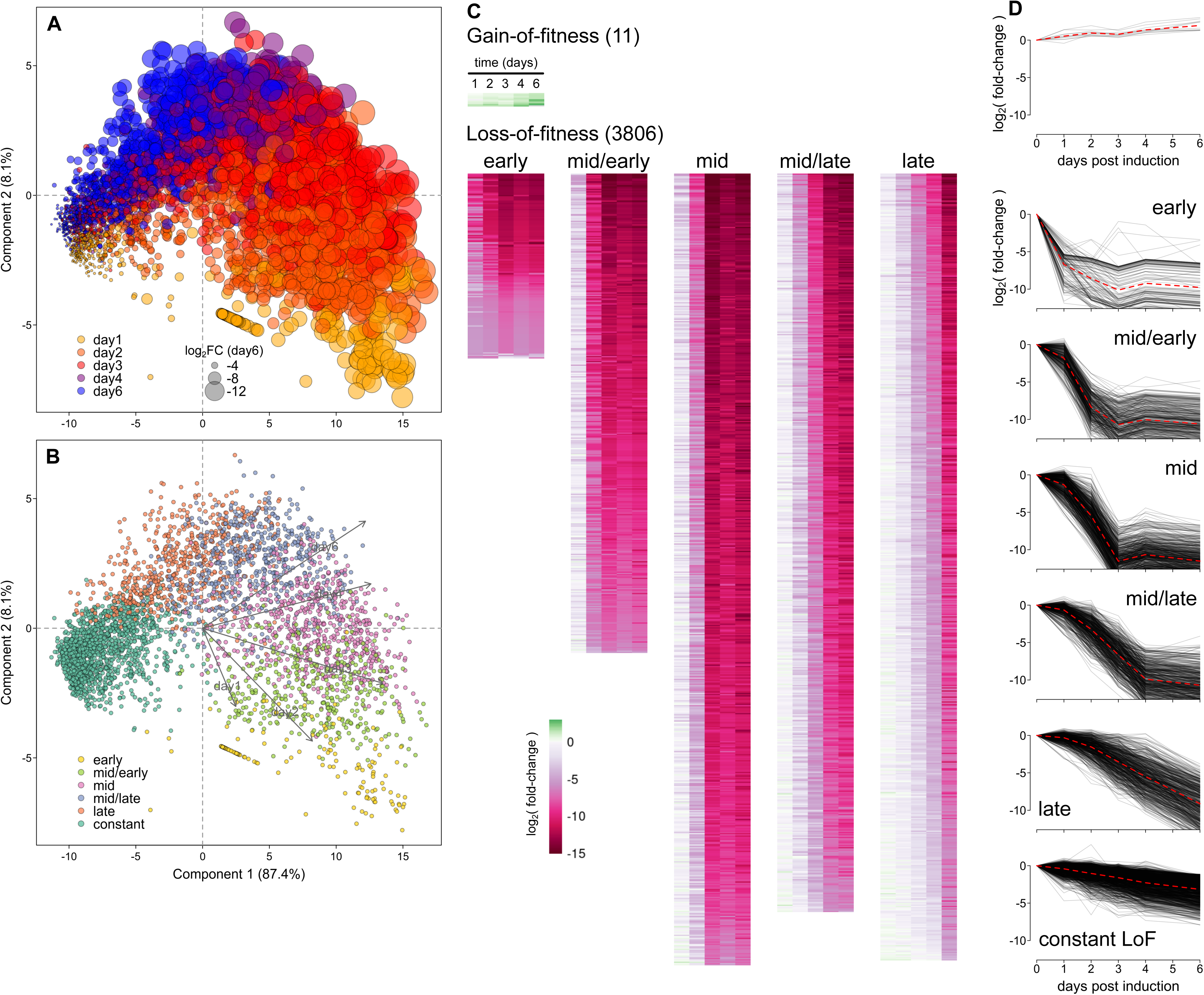
Kinetics of gene knockdown loss-of-fitness in trypanosomes. A) Principal component analysis of measured fitness cost across 6 days of RNAi induction for 3806 genes with a significant loss-of-fitness (fdr ≤ 0.01) by both total gene count Z and median fragment Z methods. Points are coloured for the day post-induction at which they show the greatest change in log fold-change (i.e. the greatest change in fitness). Circle size represents total log_2_ fold-change by day 6. B) K-means clustering of temporal effects based on temporal change in log fold-change. Clusters are mapped to the principal components shown in (A). C) Heatmaps of measured log _2_ fold-change for clusters identified in (B). D) Temporal profiles of log_2_ fold-change for genes with significant gain-of-fitness (fdr ≤ 0.01) and for 6 clusters identified in (B).

Unexpectedly, quantitative enrichment analysis of loss-of-fitness timing profiles demonstrates that many cellular components and processes have not only distinctive effect sizes (see Fig. 5A,B), but also characteristic timings for effects post-induction (Fig. 9 and Suppl. Fig. S8). Genes associated with GO terms related to the nucleosome, DNA binding and chromatin organisation are very significantly biased towards ‘early’ post-induction effects. Terms related to flagellar biology/structure arise ‘mid/early’, and targetting core components of the ribosome or those involved in ribosomal processing result in loss-of-fitness effects significantly biased towards ‘mid’ and ‘late’ time-points, respectively. This observation of GO term-specific effect timings is not merely the product of overlap between genes occurring in similar terms, as terms relating the same underlying process but with no shared components (e.g. proteasome core and regulatory particle, large and small ribosomal processome, intraciliary transport particle A and B) also occur at the same characteristic timing and enrichment (Fig. 9 and Suppl. Data File 10). Taken together, these data demonstrate that DRiF-Seq is able to provide quantitative information on both RNAi effect size and also timing associated with genes, which reveals a characteristic hierarchy to functional ablation of genes related in particular cellular structures, complexes and processes. This is not sufficiently discriminatory on its own to be able to predict gene function, but temporal fitness profile provide an important additional line of information towards the classification of genes of unknown function. It also shows that DRiF-Seq provides a means to screen for perturbation of loss-of-fitness timing in addition to effect size.

**Figure 9.**
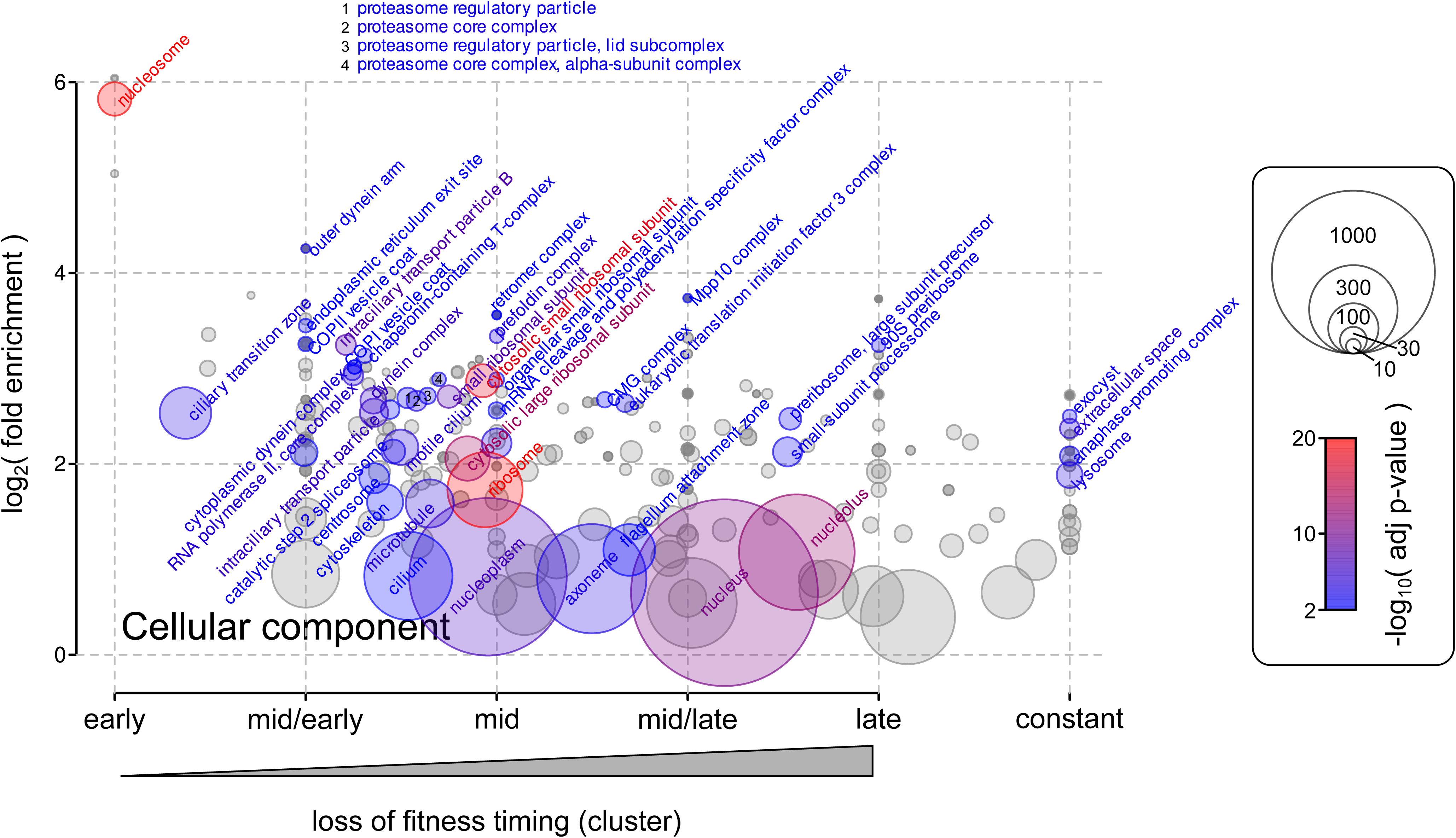
Quantitative enrichment analysis of GO terms (Cellular components) by timing of loss-of-fitness profile. GO terms that are significantly enriched (overall *p*-value ≤ 0.1) are highlighted. Circle size reflects the total number of genes encompassed by each GO term. Overall *p*-value for terms across timing profiles are the harmonic means of the adjusted *p*-values for enrichment in each of the 6 timing clusters. Enrichment and timing values are arithmetic means weighted by significance (see Methods). Data for Biological Process and Molecular Function ontologies are shown in Suppl. Fig. S8 and full data are available in Suppl. Data File 10.

## Discussion

We have described a ground-up redesign of highly-parallel RNAi experiments in African trypanosomes to produce a method we are calling Direct-RNAi Fragment Sequencing (DRiF-Seq). The method is analogous to previous genome-scale RNAi-based screens in *Trypanosoma brucei*, including RNA interference target sequencing (RIT-seq), but encompasses multiple changes including to the recipient cells, fragment library, RNAi integration site, RNAi plasmid, and sequencing of fragments. In DRiF-Seq each individual mutant in a population can be followed quantitatively through a time series, whilst knowing the precise RNAi fragment in each, allowing information about effects of targetting different parts of a gene, differential effects of fragment length and composition, and clone-clone variation to be systematically analysed.

In addition to redesign of large-scale RNAi screens, we have demonstrated *a posteriori* noise estimation (*ap*NE) as a new means to analyse these libraries – as well as other quantitative phenotypic data where all sources of variation are difficult to model or estimate through extensive replication. Combined with *ap*NE, DRiF-Seq provides quantitative fitness estimates that are sufficiently robust to identify the cost of loss of individual components from a cellular complex/process, profile fitness effects through time, and make quantitative comparisons between taxa and experimental conditions.

>50% of non-VSG genes are associated with loss-of-fitness when targetted by RNAi in EATRO1125 bloodstream form. Given the nature of RNAi in any organism, it is highly likely there will be essential genes in the genome where knock-down of RNA levels is not sufficient to see fitness effects. However, this is substantially mitigated by the ability of RNAi to target all members of gene arrays with high identity at the DNA level (which are common in trypanosomes) plus the ability to select for lethal defects in the absence of RNAi – both of which are much more challenging with knockout approaches. Moreover, the proportion of genes associated with loss-of-fitness when targetted by DRiF-Seq in *Trypanosoma brucei* is very similar to the proportion associated with loss-of-fitness in CRISPR screens in *Toxoplasma gondii* (47% (Sidik et al. 2016) and Suppl. Fig. S3) and close to knockout profiling in the highly optimized genome of *Plasmodium berghei* (63% (Bushell et al. 2017)). This is in spite of bloodstream-form trypanosomes having a heavily reduced dependence on a major cell component – the mitochondrion, which is associated with many more essential genes in insect-stages – and suggests that all of these parasites have a high proportion of fitness-associated genes compared to free-living organisms (Blomen et al. 2015; Boutros et al. 2004; Giaever et al. 2002; Kamath et al. 2003).

DRiF-Seq technology is transferable to other strains and species of trypanosome, allowing us to create a library of >200,000 RNAi mutants in a strain of *Trypanosoma brucei* which is competent for quorum-sensing and recapitulates the characteristic waves of parasitaemia associated with African trypanosome infection in the field. Not only is this the first demonstration of highly-parallel phenotypic profiling outside of the monomorphic Lister 427 line, it also offers huge potential to understand important aspects of infection biology in vivo. Coincidentally, it also revealed that RNAi is more effective in EATRO1125 than the strain used for the majority of experimental work in labs – equivalent to ∼4-fold greater effect size 3 days post-induction for essential genes (in spite of a slower doubling time for EATRO1125 in culture). The reason for this difference is unclear, but means that EATRO1125 is not only a better model for testing many aspects of trypanosome biology (including during infection), but also a more sensitive chassis for gene fitness effects. In this context, the extension of high-complexity library production to EATRO1125 is a significant contribution to trypanosome research in addition to the fitness profiling demonstrated herein. Further extension of the system to other strains, such as the most wide-spread trypanosome causing human disease *T. brucei gambiense*, is also possible and currently being developed. As is transition of the technology to the most important causative agents of animal African trypanosomiasis, *T. congolense* and *T. vivax*. Importantly, quantification of fitness effects using DRiF-Seq were sufficiently robust to identify aspects of the biology that are differentially sensitive even between 2 closely trypanosome strains – effects that were reproducible with individual RNAi lines and predictive of differential drug sensitivity. This robustness and sensitivity means that extension of the method to other species will allow use of genome-wide screens to address specific aspects of disease where quantitation of effects is critical (for example, in understanding tissue tropism or sensitisation to drug) and also to understand systematic differences in the fundamental biology of the species.

In addition to the loss-of-fitness effect size for individual gene knockdowns, DRiF-Seq reveals that the timing of effects is also informative. We have classified 14,211 trypanosome genes into 8 fitness profiles (including no change in fitness). Genes associated with specific cellular components and processes have not only distinctive effect sizes, but also characteristic timings for when these effects become apparent following induction. The effect is surprisingly precise, with independent parts of complexes such as the ribosome or proteasome having very similar measured phenotypic timings. Although unexpected, this makes some sense in the context of the organism biology, as compromising a complex or pathway by removing different essential components is likely to have similar phenotype in the cell, but does imply a surprising degree of correlation of protein levels and turnover characteristics for genes that contribute to specific processes. Kinetic fingerprints are certainly not sufficient to be able to place unknown components into specific cellular locations/processes, but may be used to exclude likely gene involvement in certain processes or distinguish possible sites of action for proteins localising to multiple parts of the cell.

There are only 11 genes associated with significant gain-of-fitness when knocked-down in EATRO1125 cultures (compared to 3806 associated with loss-of-fitness under the same criteria). These include subunits of 5’-AMP-activated protein kinase, which has been implicated in transition from proliferation to quiescence (Saldivia et al. 2016), a tRNA modifying enzyme and several proteins of unknown function, but share no common components with GoF genes in Lister 427. As fitness was deliberately assessed under culture conditions in which density-dependent quorum-sensing would not be triggered, the sets from neither strain include genes previously identified in RNAi screens looking for quorum-sensing factors.

In summary, DRiF-Seq provides temporal fitness landscapes for individual genes at a clone-by-clone level and also specific processes. It appears to be highly robust and sensitive for quantitative fitness cost measurements enabling comparison both between strains and across large phylogenetic distances in comparative functional genomic analyses between very different parasites. The scale and resolution of the method will be applicable to multiple situations and there is great potential for further extension in analysis of epistatic interactions or phenotyping in host.

## Materials and methods

### Cell culture and transformation

Bloodstream form *Trypanosoma brucei* strain Lister 427 (expressing VSG-2/MITat1.2) and strain EATRO1125 (expressing VSG AnTat1.1E) plus all derivatives of these strains were grown in HMI-9 medium supplemented with 15% foetal bovine serum at 37°C and 5% CO_2_ (Hirumi and Hirumi 1989).

Routine transfection of individual plasmids was as in (Schumann Burkard et al. 2011). Briefly, 2-3×10^7^ cells were harvested from actively-dividing cultures (<1×10^6^ cells ml^-1^ for EATRO1125), resuspended in 120 µl 90 mM sodium phosphate, 5 mM KCl, 0.15 mM CaCl_2_, 50 mM HEPES, pH 7.3 containing 10-20 μg of linearised DNA, and electroporated in 2 mm cuvettes using an Amaxa Nucleofector 2b device (Lonza) with program Z-001. Cells were transferred to flasks containing pre-warmed medium and allowed to recover for 8-16 h, after which antibiotic selection was applied. Antibiotic concentrations were 0.1 μg ml^-1^ puromycin (pSmOx), 5 μg ml^-1^ hygromycin B (pV4-Sce and derivatives) or 2 μg ml^-1^ phleomycin (p2T7 derivatives). Where used, induction of RNAi was by addition of 1 μg ml^-1^ tetracycline to culture medium following selection of stable transformant lines.

To estimate transfection efficiency, samples of cells immediately following recovery were diluted in fresh, selective medium and distributed across 96-well plates. For routine transfections 20% and 4% of each transfection was plated on 2 independent plates (providing estimates of efficiency with <30% relative error in the range 200-8000 independent clones). Numbers of independent transfectants were derived from the number of positive wells based on Poisson-distributed filling of wells. Confidence intervals of estimates were derived from the distribution of quantiles of 10^6^ simulated platings at 0.001 to 10 cells well^-1^.

### Generation of DRiF-library recipient cells

To generate a self-excising construct that creates a double-strand break (DSB) at a silent minichromosomal VSG, a construct was designed to target sequence encoding I-SceI with an N-terminal nuclear localisation signal from pLew100::NLS-ISceI-HA (Boothroyd et al. 2009) to *VSG427-31* (*VSG-G4*) in the same arrangement as used for high-efficiency transfection of *T. congolense* (Awuah-Mensah et al. 2021). Expression of NLS-I-SceI is controlled by a tetracycline-inducible *T. brucei* rRNA promoter independent of selection marker. An I-SceI recognition site upstream of the NLS-I-SceI cassette generates a DSB on induction and subsequent loss of downstream (telomeric) sequence in the absence of recombination. This construct, pV4-Sce, was transfected into Lister 427 SmOx cells as described above. Clones were tested for loss of *HYG* and also transfection efficiency in the presence and absence of induction of NLS-I-SceI expression, with no evidence of clone-clone differences or instability on freeze-thaw (Suppl. Fig. S1). A single clonal line was therefore taken forward as 427-TTS (for <u>T</u>7 RNA polymerase, <u>T</u>etR, and I-<u>S</u>ce-I).

For *T. brucei* strain EATRO1125, cells harvested from a mouse infection were first modified with the Single Marker Oxford construct, pSmOx (Poon et al. 2012) to create a line herein called AnTat-SmOx. RNA-Seq confirmed that this line was still expressing VSG AnTat1.1E (in addition to expected transgenes) and competency for inducible RNAi was confirmed by transfection with p2T7-177-CHC (Sarah Whipple, University of Nottingham) targetting the clathrin heavy chain gene (*CHC*), which causes a well-characterised enlarged flagellar pocket phenotype. pV4-Sce was modified to target the same payload above but also the *VSG427-31* integration sites to the EATRO1125 minichromosomal *VSG* KX698662 (pAnTat-V1-V4-Sce). Transfection into AnTat-SmOx created a line, AnTat-TTS, in which an induced DSB in EATRO1125 can be targetted with plasmids integrating at the 427-specific *VSG427-31*.

### Generation of DRiF-Seq plasmid libraries

For DRiF-library production, a specific version of the inducible RNAi vector p2T7 (LaCount et al. 2000) was generated. p2T7-V4δ targets the RNAi and selection cassettes of p2T7 to *VSG427-31* without the parts of the plasmid necessary for bacterial growth and contains a modified fragment cloning site compatible with dual 8-cutter cloning (SfiI-SbfI). A NotI site in the pre-loaded stuffer prevents carry-over of unmodified plasmid into cells.

To generate genome-wide targeting fragments, 20 μg strain-specific gDNA was sonicated in SSC buffer (150 mM NaCl, 15 mM sodium citrate, pH7.0) on ice at 25% intensity, 40% of the time for 4 minutes (Bandelin SonoPuls). Sheared DNA was concentrated by isopropanol precipitation and fragments of approximately 400-500 bp were size selected from an agarose gel. Size-selected fragments were end repaired (NEBNext End Repair Module) at 20°C for 2 hours. Directional fragment-end adapters were generated by annealing custom primers (see Suppl. Data 11) and added to 500 ng of fragments by ligation at 20x molar excess with 10u T4 DNA ligase at 16°C for 2 hours. To improve ratio of amplifiable fragments with different adapters at each end a ‘vectorette’ approach was taken (Arnold and Hodgson 1991) with a 4:1 molar ratio between vectorette and conventional adapter. Linked fragments were separated from unattached adapter by gel electrophoresis, amplified using primers 28s and 2fr (Suppl. Data 11), digested with SfiI and SbfI and ligated into 250 ng p2T7-V4δ cut with SfiI and NsiI at a 4-fold molar excess at 16°C for 4 hours. Reaction was desalted by silica-column purification (Monarch Spin PCR & DNA Cleanup Kit; NEB) and ∼100 ng used to transform NEB10-β Competent *E. coli* by electroporation at 20 kV cm^-1^ and 25 μF. Transformed libraries were sampled to measure complexity and the remainder grown directly in 200 ml Luria Broth medium supplemented with 100 µg ml^-1^ ampicillin, followed by purification of plasmid DNA by anion exchange chromatography (Qiagen Plasmid Maxi Kit). Plasmid libraries represented 7.6×10^6^ and 2.7×10^6^ independent fragments for Lister 427 and EATRO1125 strain libraries, respectively.

### Generation of DRiF-Seq libraries in trypanosomes

For transfection of Lister 427 cells, selective pressure for integration site (hygromycin B) was removed from 427-TTS cells for 48 h followed by induction of DSB for 16 h with 1 µg ml^-1^ tetracycline. A total of 5.4×10^8^ cells were harvested from actively-dividing cultures by centrifugation, washed to remove residual tetracycline and then resuspended in 0.9 ml Tb-BSF containing 135 µg NotI-cut plasmid library. This was distributed across 9 transfection cuvettes (each containing 60×10^6^ cells and 15 µg cut plasmid), each of which was electroporated on a Amaxa Nucleofector 2b device (Lonza) using program ‘Z-001’, as above. Transfections were mixed and then immediately split in half to create 2 libraries of independent RNAi mutants. Samples of 1/2000 and 1/10000 of the total were distributed across 96 well plates to estimate library complexity and the remainder selected as populations using 2 µg ml^-1^ phleomycin. Lister 427 DRiF-Seq libraries had a total estimated complexity of 6.1×10^5^ independent clones (4.7 -7.7×10^5^ 90% confidence interval) split across 2 independent libraries. Libraries were maintained at > 20x complexity to avoid loss of complexity.

EATRO1125-strain AnTat-TTS cells were transfected similarly to 427-TTS cells except that a total of 2×10^9^ cells distributed across 25 cuvettes (80×10^6^ cells in each) was used, and samples of 1/800 and 1/2400 of the total were used to estimate library complexity (∼2.5×10^5^ independent clones; 2.1 -3.1×10^5^ 90% confidence interval). AnTat-TTS libraries were maintained at ≤ 1×10^6^ ml^-1^ during selection to prevent pressure from quorum sensing.

### Sequencing of DRiF-Seq libraries

For quantitative analysis of mutant abundance, cells were grown such that no bottleneck smaller than 250x library complexity was encountered. RNAi was induced by addition of 1 µg ml^-1^ tetracycline and all samples were made in duplicate at > 120x complexity. Cells were harvested by centrifugation, washed twice in TNEP (120 mM NaCl, 20 mM Tris-HCl, 14 mM Na_2_HPO_4_, 1.5 mM KH_2_PO_4_, 10 mM EDTA, pH8) and resuspended in 200 µl TNEP containing 200 µg ml^-1^ RNase A before isolation of genomic DNA by binding to silica (DNeasy Blood and Tissue kit, Qiagen). Care was taken that dilution from growth and washing would be sufficient to reduce any residual non-integrated plasmid remaining from library transfection to ≪ 1 copy per fragment and that DNA from the total sample size could be captured at near 100% yield without saturation of the column. DNA was eluted in 10 mM Tris-HCl, pH8.5.

1% of each sample was taken for qPCR against purified plasmid standards to check that isolated RNAi fragment copy number matched input sample cell count, and RNAi fragments from the entire remaining DNA were then amplified by linear PCR in 300 µl reactions using primers 2fw and 137 (Suppl. Data 11). Standard Taq PCR conditions were used with the addition of 0.5 mM EvaGreen plus (Biotium), 1 mg ml^-1^ BSA and 5% (v/v) dimethyl sulphoxide. PCRs were stopped while amplification was still exponential and at the same degree of amplification for all samples. Amplicons were purified by silica-binding (PCR purification kit, Qiagen) and a low-cycle, linear nested PCR used to add standard Illumina flow-cell binding sequences (but not sequencing adapters) and sample indices to amplicons (Suppl. Data 11). A single, large scale isolation and indexing PCR can be used in place of this approach, but nesting provides additional convenience for the large PCRs as well as choice of index combination for sequencing.

Fragment-length distributions were analysed using a TapeStation 4200 and High Sensitivity D1000 ScreenTape Assay (Agilent). Libraries were pooled in equimolar amounts and final library quantification performed using the KAPA Library Quantification Kit for Illumina (Roche). Indexed pools were sequenced using a NextSeq 500 System (Illumina) with high-output 150 cycle kit v2.5 (Illumina). A minimum of 40M paired-end reads passing filter per timepoint per sample were generated, providing sequencing coverage at > 130x and > 200x library complexity for Lister 427 and EATRO1125-based libraries, respectively.

### Sequence data processing

Raw sequence data were demultiplexed to individual samples with bcl2fastq2 (v2.20) with no index mismatches allowed. The first 2 bases of Read2, which are common to all fragments, were removed with fastx_trimmer (v0.0.14) from FASTX-Toolkit (github.com/agordon/fastx_toolkit) and reads were trimmed for quality with trim_galore (v0.6.10) in paired mode retaining only pairs with at least 30 base of sequence at each end passing filter (-q 28 --paired --stringency 20 -r1 30 -r2 30). Trimmed reads were mapped to the *T. brucei* Lister 427 long-read plus Hi-C assembly available at tritrypdb.org v48 (Alvarez-Jarreta et al. 2024) with bowtie2 v2.5.2 (Langmead and Salzberg 2012; Langmead et al. 2019), allowing fragments of length 150-750 bp (--no-mixed --no-discordant -I 150 -X 750). Counts of mapping fragments starting and ending at exactly the same reference position and in the same orientation were extracted with a custom script with an overall fragment MAPQ calculated from the mean of all contributing read pairs. Only fragments with counts ≥ 24 across all time-points were taken forward for analysis.

### Analysis of fitness effects

To compare change in individual fragment counts in the population following induction of RNAi, raw count data were converted to Bland-Altman coordinates using the standard transformations *A* = 0.5*log_2_((*c_0_*+0.5)·(*c_t_*+0.5)), *M* = log_2_((*c_t_*+0.5)/(*c_0_*+0.5)), where *c_0_* and *c_t_*are the read counts at times 0 and *t* post-induction, respectively. Assuming that most fragments (which include ∼60% non-genic DNA) cause no fitness change and sources of variance will sum to an approximately Gaussian dispersion in an abundance-dependent manner, the data were separated into 24 abundance (*A*) quantiles and spread of top 75% of *M* for each quantile fitted to a censured Gaussian by non-linear least-squares fitting (Gauss-Newton algorithm, initial estimates taken from the data). Fitting to a censured Gaussian prevents underestimate of dispersion due to exclusion of data and does not prescribe the amount of data having either loss- or gain-of-fitness. Fits to the data strongly support the validity of these assumptions (Fig. 2A,B). The mean and variance of the fitted Gaussians were then used to fit abundance-dependent parameter models (see Fig. 2C), which were used to infer Z scores for fragment changes. P-values derived from Z scores were adjusted for multiple comparisons by the Benjamini-Hochberg method.

Gene-wise effects were assessed using 2 metrics derived from fragment-wise measurements. Median Z scores were calculated based on individual fragments having ≥ 100 bp overlap with an annotated gene. Alternatively, summed read counts for all fragments overlapping each gene were used to derive gene-wise Z scores using the same *a priori* noise-estimation approach as used for individual fragments (except that data were binned into 16 abundance quantiles). Median fragment Z scores capture better the behaviour of the average clone containing an RNAi fragment targeting a specific gene, while sum gene-wise Z scores reflect the overall change in abundance for all reads derived from each gene, but the 2 metrics show very good agreement (see Fig. 2F). ‘Mixed’ fragments overlapping more than one annotated gene were excluded from analysis.

### Temporal analysis of fitness cost profiles

A stringent set of 3806 genes associated with loss-of-fitness when targetted in DRiF-Seq libraries was determined based on genes having fdr ≤ 0.01 from both Z score of change in total gene count and median Z of all contributing fragments. 11 genes are associated with significant gain-of-fitness under these criteria. Principal components were calculated from the estimated overall gene-wise log_2_ fold-change against day 0 at the 5 experimental sample days (days 1,2,3,4,6 post-induction). Temporal clustering was by the *k-*means method using the change in effect size (Δlog_2_ fold-change) between samples in the time-series. Optimal number of clusters (*k*=6) was calculated using the gap statistic method (Tibshirani et al. 2001) applied by the R package NbClust (Charrad et al. 2014).

### Quantitative enrichment analysis

For quantitative enrichment analysis of fitness effects in *T. brucei*, Lister 427 genes were converted to TREU927 identifiers using orthologs identified by TriTrypDB v58 (Alvarez-Jarreta et al. 2024) and these were then mapped to GO terms annotated for TREU927 (v58). A total of 7420 genes with measured fitness have GO term annotation in TREU927, covering 63717 annotations of 3453 unique GO terms. Phenotypic scores for each gene were mapped to every de-duplicated GO term associated with that gene, providing a set of scores for each observed term made up of the scores for each observed gene contributing to it. Only GO terms with scores from at least 3 genes were analysed. Distributions of set scores were tested for being more extreme than expected from the population distribution by Monte Carlo permutation test (10^7^ replicates), with Benjamini-Hochberg correction for multiple hypothesis tests.

To gain an overall adjusted p-value for enrichment of terms across timing profiles, values for enrichment of GO terms in each of the 6 timing clusters (‘early’ to ‘late’, plus ‘constant’) were combined as harmonic mean *p*-values as a means to control false positive rate across mutually exclusive, but not independent tests (Wilson 2019). Enrichment and timings are the arithmetic means across the clusters, weighted by significance of enrichment ({-log_10_(adjusted *p*-value)}^-2^).

### Comparison of fitness costs between parasite species

Comparisons were made between measured fitness costs in our DRiF-Seq EATRO1125 library (log_2_ fold-change at day 6 post-induction), *Toxoplasma gondii* “phenotype” (mean log_2_ fold change for the 5 guides with highest effect at passage 6; (Sidik et al. 2016)), and relative growth rates of *Plasmodium berghei* gene knockouts (Bushell et al. 2017). Genes in *T. brucei* Lister 427 were converted to TREU927 identifiers according to the orthologs identified by TriTrypDB v58 (Alvarez-Jarreta et al. 2024) and genes were then mapped to ortholog groups based on their classification in OrthoMCL-DB v6 (Chen et al. 2006). A total of 8132, 8158 and 2406 genes (*Trypanosoma*, *Toxoplasma*, and *Plasmodium*, respectively) have available fitness data from genome-scale phenotyping and are also mapped to orthoMCL-DB groups, covering 6984, 7420 and 2338 ortholog groups. 761 ortholog groups have gene fitness data available for all 3 organisms.

To bring all observations into the range 0-1, fitness costs were normalised based on the range of observed scores within each set. For binary comparison of ortholog group loss-of-fitness (Fig. 7B), loss-of-fitness in an organism was defined as a normalised fitness cost of ≥ 0.3. GO terms for genes were taken from the respective organism datasets in TriTrypDB (v58), ToxoDB and PlasmoDB (v51). To mitigate bias in GO terms resulting from different types of annotation datasets being available for each of the organisms, GO terms associated with any gene contributing to an ortholog group were mapped to that orthoMCL group. Quantitative enrichment analysis was performed as above.

### Cell growth rate analysis for individual RNAi mutants

To quantify growth of individual RNAi lines, an ATP-based luciferase assay was used. Cells were diluted to 10^3^ cells ml^-1^ in 1.5 ml wells on a 96 deep-well plate and sampled at 24-hour intervals. Prior to sampling, wells were mixed and 100 μl culture transferred to black 96 well plates. 50 μl of CellTiter-Glo Reagent 2.0 (Promega) was added to cells, incubated for 10 min, and luminescence detected using a Glomax 96 Microplate Luminometer (Promega). A cell standard series was used at the start of the experiment to establish linear range and sensitivity, and a set of ATP standards covering the linear range included on each plate to normalise plate signal between time points. Cultures were diluted depending on cell density to maintain below quorum sensing limits.

### Infections with bloodstream-form T. brucei

Male C57BL/6J mice (8 weeks old) were infected intraperitoneally with 2,000 parasites and peripheral parasitaemia was monitored daily by venipuncture of the lateral tail vein followed by haemocytometry.

### Ethics statement

Animal experiments were performed under UK Home Office regulations (project licence P8E71BFCF) and Portugal Direção Geral de Alimentação e Veterinária (project licence 018889\2016). Research was ethically approved by the University of Nottingham Animal Welfare and Ethical Review Board and Órgão Responsável pelo Bem-estar Animal (ORBEA) of Instituto de Medicina Molecular.

## Acknowledgements

We are grateful to Nadine Holmes in the Deep Seq facility, University of Nottingham, for Illumina sequencing. This work was supported by University of Nottingham/Wellcome Trust Institutional Strategic Support Fund 204843/Z/16/Z award to BW; MRC New Investigator Award MR/N01037X/1 to CG; and NC3Rs Project Grant NC/W001144/1 to BW and CG.

