## Supplemental Figures for "A platform for robust quantitative functional genomics reveals temporal fitness landscapes in African trypanosomes"

#### Supplemental figure legends

**Supplemental figure S1**. High efficiency transfection in DRiF-library recipient cell lines 427-TTS and AnTat-TTS. A) Consistency and stability of high transfection efficiencies across clones containing a DRiF-compatible self-cleaving construct inserted at a silent *VSG* locus on a minichromosome. Double-strand break at the target locus is induced by the addition of tetracycline (+Tet). Optimisation of time of double-strand break induction (B) and mass of plasmid DNA (C) in 427-TTS. D) Performance of a similar strategy in EATRO1125 bloodstream-form cells.

**Supplemental figure S2**. Comparison of measured gene effect sizes and variance in RIT-Seq and DRiF-Seq experiments in Lister 427 and EATRO1125 at different times post-induction of RNAi. RIT-Seq data are from 7435 non-redundant TREU927 genes in *(Alsford et al. 2011)*. Genes identified as associated with a significant gain- or loss-of-fitness in previous analysis by DEGSeq ($|Z| > 3.3$) are indicated (red) alongside reanalysis of the data by *a posteriori* noise estimation (blue lines at fdr = 0.1 and 0.01). DRiF-Seq libraries in Lister 427 (16152 genes) or EATRO1125 (14211 genes) show *ap*NE fdr only. Gene sets associated with select biological functions are highlighted as in Fig. 4.

**Supplemental figure S3**. Application of *a posteriori* noise estimation to genome-wide CRISPR data from *Toxoplasma* (Sidik et al., 2016). A) Calculation of Z scores for 8158 targetted genes based on either the sum of counts for all sgRNAs targetting a gene or the median sgRNA effect for each gene. The expected null distribution estimated from the fitted variance of the data is overlaid in red. The number of genes detected by *ap*NE as enriched/depleted at fdr = 0.1 or 0.01 are shown in blue, in addition to percentage over-representation of genes in each distribution tail. B) Distribution of 'phenotype' scores (mean $\log_2$ fold change in the normalised counts of the top five scoring sgRNAs for each gene) showing the position of minimum Z-scores (genes inferred by *ap*NE to have no effect). C) Volcano plot showing distribution of adjusted p-values for genes calculated by *ap*NE against their P6 phenotype score.

**Supplemental figure S4**. Quantitative enrichment analysis for fitness effect sizes associated with GO Terms in Biological Process, Cellular Component and Molecular Function ontologies. Terms that have fitness effect sizes significantly different (adjusted p-value ≤ 0.05; Monte Carlo permutation test) from the average effect size of genes with GO annotation. Circle size indicates number of measured genes associated with each GO term. For clarity, GO descriptors for some terms with lower significance are not shown. Full data are available in Suppl. Data File 8.

**Supplemental figure S5**. Distribution of DRiF-Seq fitness effect sizes for gene products localised to different subcellular compartments. Localisation annotation is from the TrypTag genome-wide protein localisation database *(Billington et al. 2023)*. Effect sizes are for EATRO1125 at 6 days post-induction. The number of genes associated with each annotation and the proportion with significant fitness cost at fdr ≤ 0.01 are indicated above each distribution.

**Supplemental figure S6**. Comparison of mRNA and protein characteristics with DRiF-Seq fragment fitness cost. All effect sizes are for EATRO1125 at day 6 post-induction. A) Mean effect size distributions for fragments targetting annotated 5'UTR, CDS or 3'UTR of 123 genes with fragments specifically targetting all 3 regions. Data are derived from a remapping of the fragment count data to the TREU927 genome to take advantage of the UTR annotation and only fragments entirely encompassed within each feature are included. B) Relationship between differential fragment effect size (difference between specific fragment effect size and median effect of targetting fragments) and differential GC content of fragment. Fragments targetting 1280 genes associated with large loss-of-fitness (≥ 256-fold depletion; fdr ≤ 0.01) and targetted by ≥ 2 fragments are included. C) Relationship between measured mRNA *(Fadda et al. 2014)* and protein *(Tinti et al. 2019)* turnover in bloodstream-form cells and DRiF-Seq effect size. Correlations for both all genes for which data are available (All) and only those with ≥ 256-fold depletion at fdr ≤ 0.01 (LoF) are shown. Box-whisker plots for 8 quantiles are also shown. D) Proportion of genes with 'hypothetical', 'putative' or full functional annotation in each of 8 quantiles of mRNA half-life. E) Distributions of measured mRNA or protein half-lives for genes with different loss-of-fitness temporal classification (Fig. 8). Circle size above each category shows the number of genes for which data are available.

**Supplemental figure S7**. Validation of differential RNAi effect size inferred from genome-wide DRiF-Seq fitness estimates by individual RNAi. Growth of 2 clonal RNAi lines for RNAi targetting 10 genes in both Lister 427 (L427) and EATRO1125 (AnTat) in the presence and absence of induction of RNAi by tetracycline are shown. For clarity, cumulative cell numbers of Lister 427 cells have been offset by 100-fold. Arrows indicate times during growth where cultures were passaged into fresh medium.

**Supplemental figure S8**. Quantitative enrichment analysis for timing of loss-of-fitness profiles of GO Terms in Biological Process, Cellular Component and Molecular Function ontologies. GO terms that are significantly enriched (overall $p$-value ≤ 0.1) are highlighted. Circle size reflects the total number of genes encompassed by each GO term. Overall $p$-value for terms across timing profiles are the harmonic means of the adjusted $p$-values for enrichment in each of the 6 timing clusters. Enrichment and timing values are arithmetic means weighted by significance (see Methods). Descriptors are given for terms with significant enrichment. Full data are available in Suppl. Data File 10.

### Suppl. Fig. 1

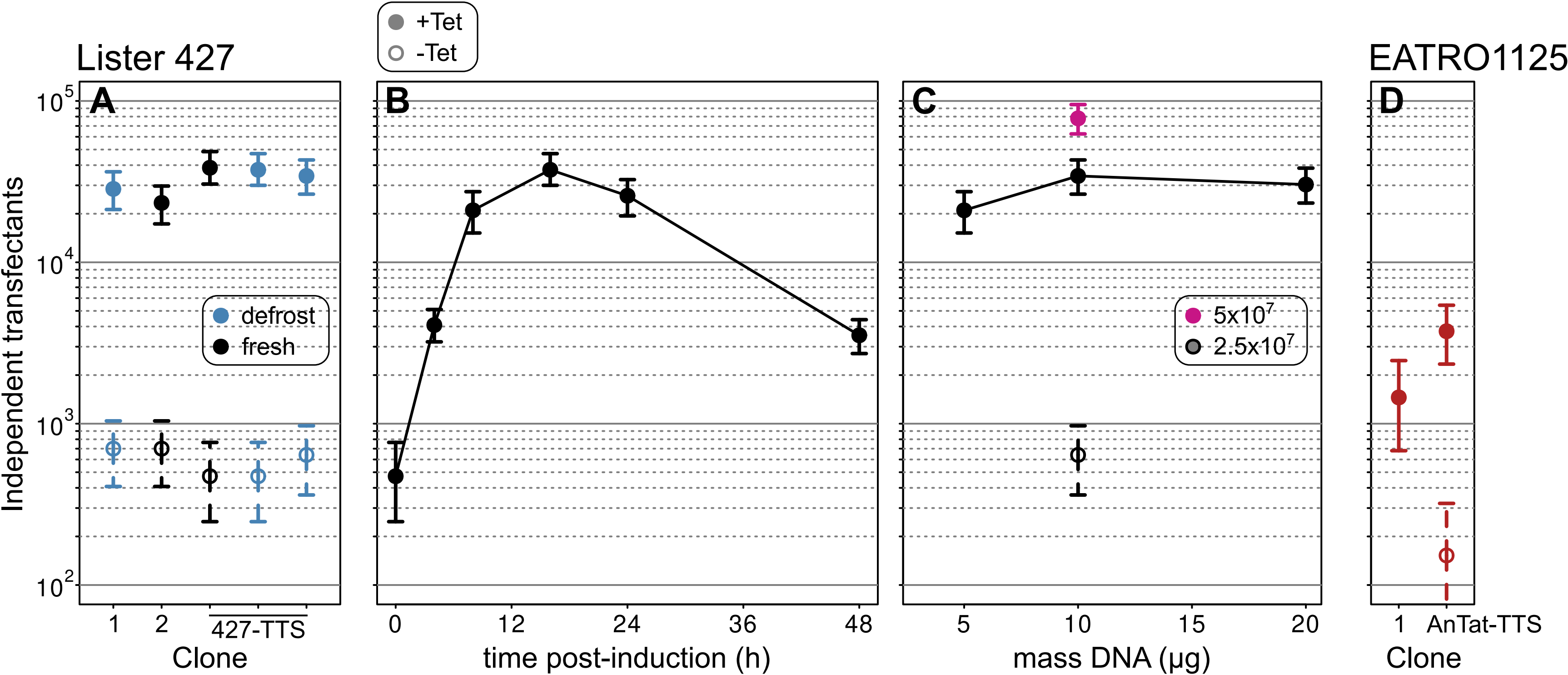
+Tet
-Tet
Lister 427
EATRO1125
A
B
C
D
Independent transfectants
10^5
10^4
10^3
10^2
defrost
fresh
5x10^7
2.5x10^7
1
2
427-TTS
Clone
0
12
24
36
48
time post-induction (h)
5
10
15
20
mass DNA (µg)
1
AnTat-TTS
Clone

Suppl. Fig. 2

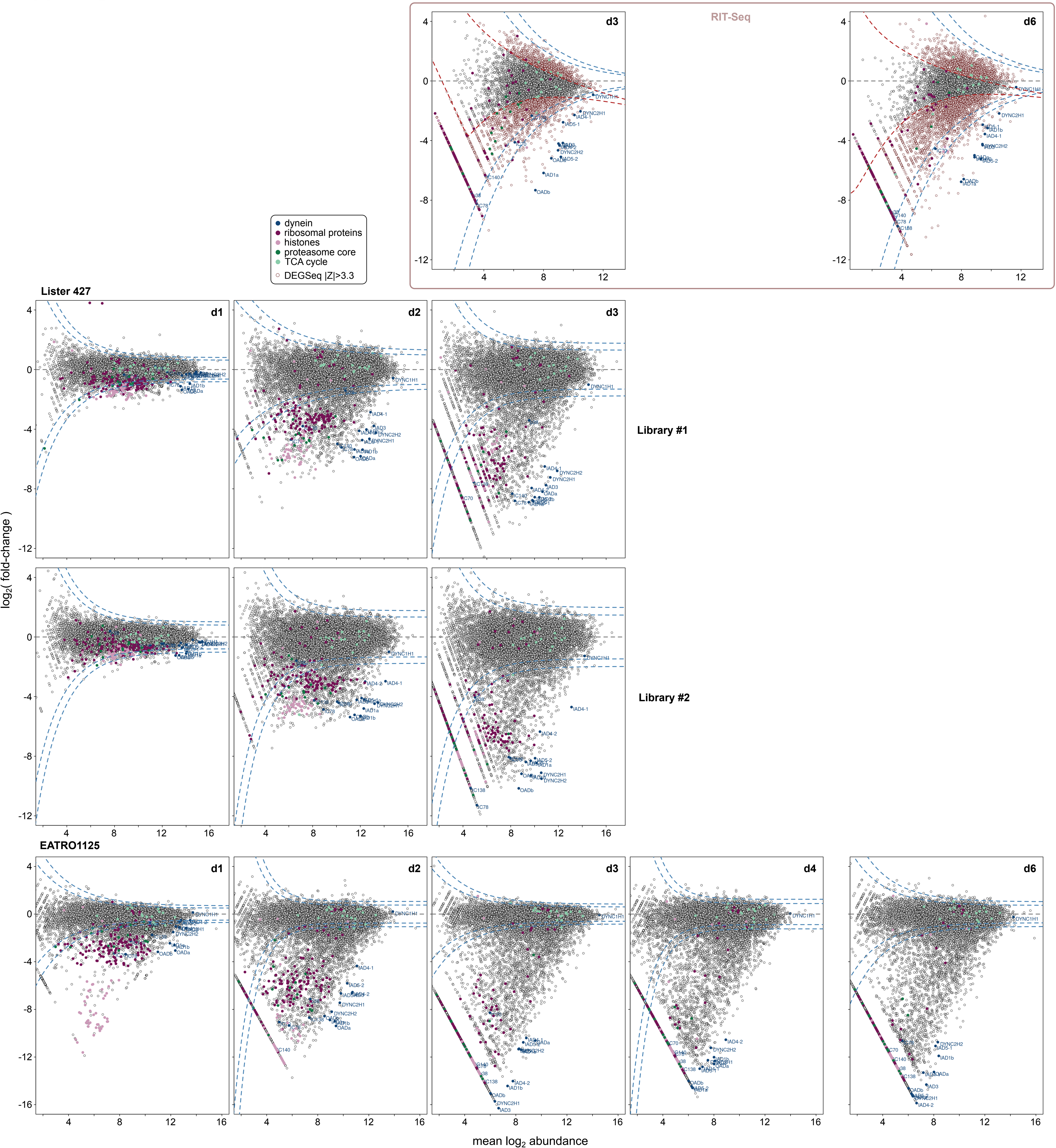
RIT-Seq
d3
d6
dynein
ribosomal proteins
histones
proteasome core
TCA cycle
DEGSeq |Z|>3.3
Lister 427
d1
d2
d3
Library #1
Library #2
EATRO1125
d1
d2
d3
d4
d6
log2( fold-change )
mean log2 abundance

### Suppl. Fig. 3

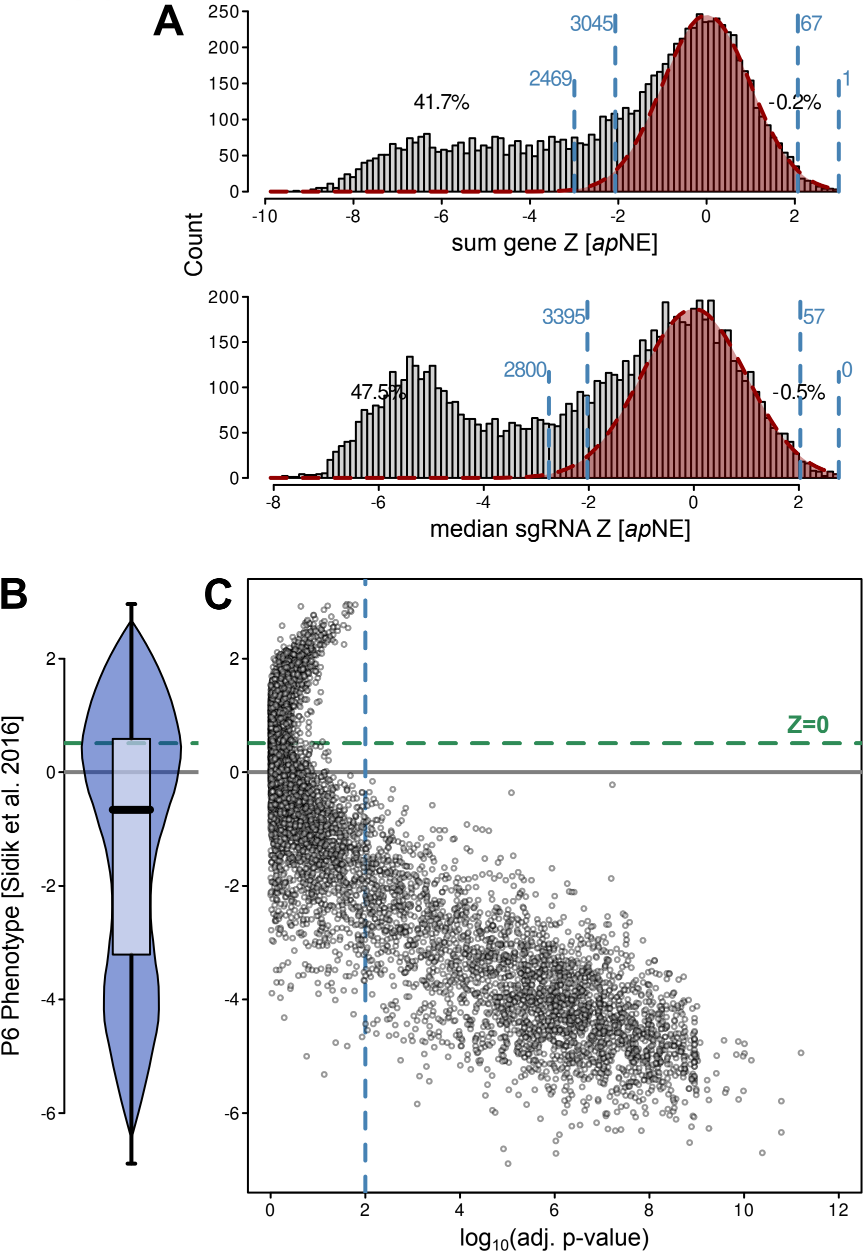
A
Count
sum gene Z [apNE]
3045
2469
41.7%
67
1
-0.2%
median sgRNA Z [apNE]
3395
2800
47.5%
57
0
-0.5%
B
P6 Phenotype [Sidik et al. 2016]
C
Z=0
log10(adj. p-value)

Suppl. Fig. 4

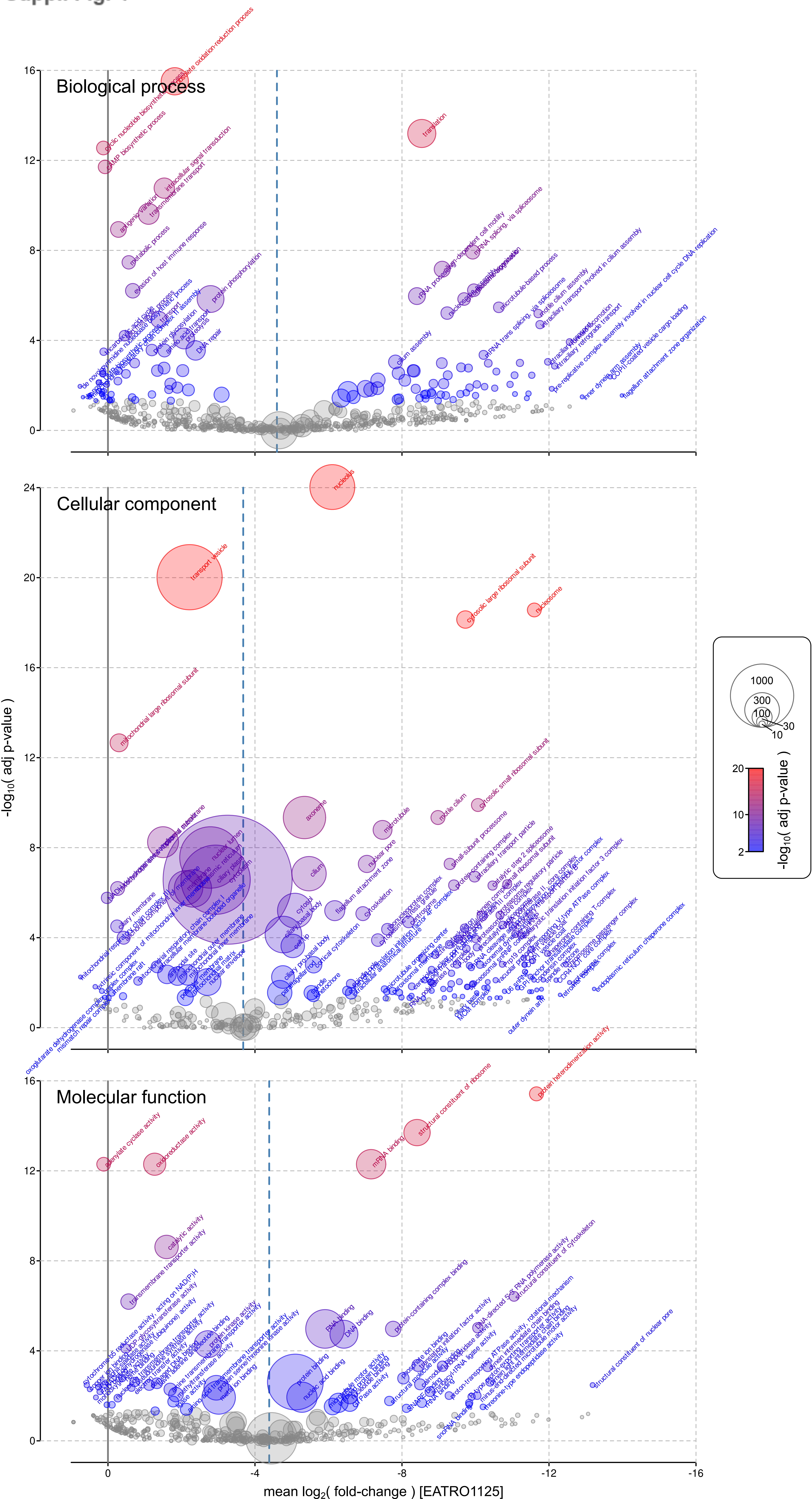

Biological process
translation
cyclic nucleotide biosynthetic process
cAMP biosynthetic process
intracellular signal transduction
transmembrane transport
antigenic variation
metabolic process
evasion of host immune response
protein phosphorylation
DNA repair
cilium assembly
rRNA processing
mRNA splicing, via spliceosome
microtubule-based process
Cellular component
nucleolus
transport vesicle
cytosolic large ribosomal subunit
nucleosome
mitochondrial large ribosomal subunit
cytosolic small ribosomal subunit
axoneme
microtubule
motile cilium
cilium
nuclear pore
cytoskeleton
Molecular function
protein heterodimerization activity
structural constituent of ribosome
mRNA binding
adenylate cyclase activity
oxidoreductase activity
catalytic activity
transmembrane transporter activity
RNA binding
DNA binding
protein-containing complex binding
structural constituent of cytoskeleton
structural constituent of nuclear pore
protein binding
nucleic acid binding
-log10( adj p-value )
mean log2( fold-change ) [EATRO1125]
1000
300
100
30
10

Suppl. Fig. 5

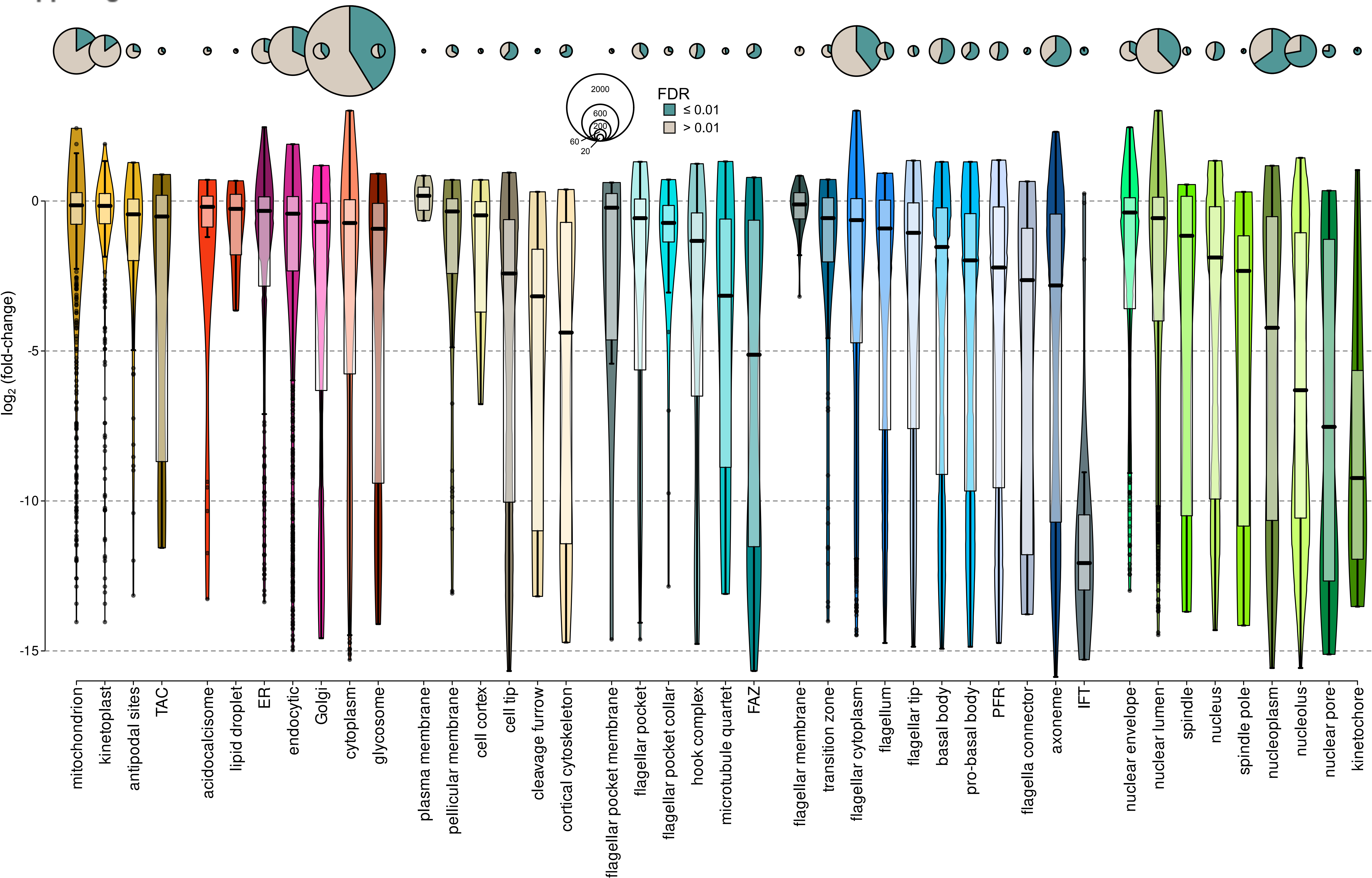

2000
600
200
60
20
FDR
≤ 0.01
> 0.01
log2 (fold-change)
0
-5
-10
-15
mitochondrion
kinetoplast
antipodal sites
TAC
acidocalcisome
lipid droplet
ER
endocytic
Golgi
cytoplasm
glycosome
plasma membrane
pellicular membrane
cell cortex
cell tip
cleavage furrow
cortical cytoskeleton
flagellar pocket membrane
flagellar pocket
flagellar pocket collar
hook complex
microtubule quartet
FAZ
flagellar membrane
transition zone
flagellar cytoplasm
flagellum
flagellar tip
basal body
pro-basal body
PFR
flagella connector
axoneme
IFT
nuclear envelope
nuclear lumen
spindle
nucleus
spindle pole
nucleoplasm
nucleolus
nuclear pore
kinetochore

### Suppl. Fig. 6

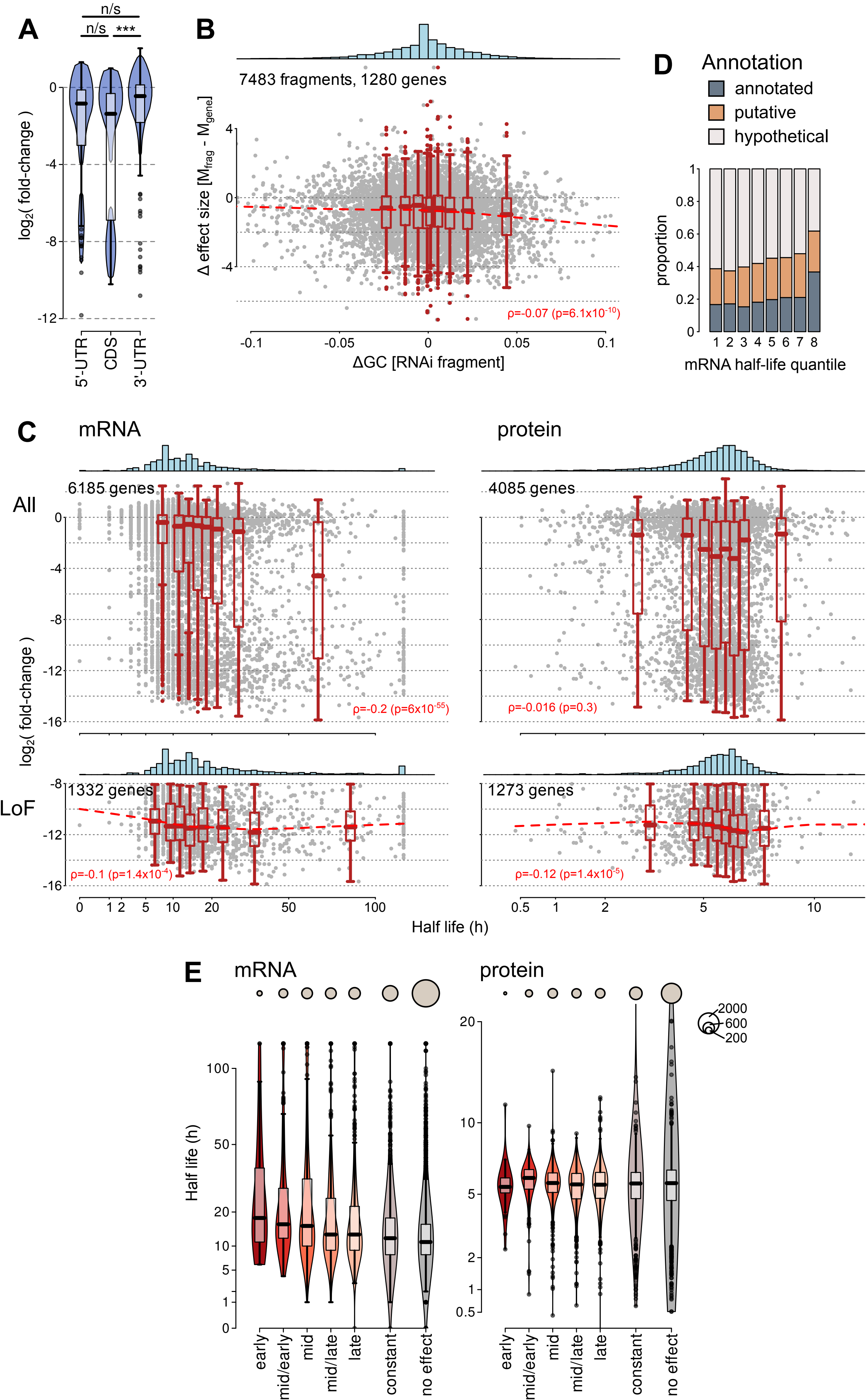
A
n/s
n/s
***
log2( fold-change )
5'-UTR
CDS
3'-UTR
B
7483 fragments, 1280 genes
Δ effect size [Mfrag - Mgene]
ρ=-0.07 (p=6.1x10-10)
ΔGC [RNAi fragment]
D
Annotation
annotated
putative
hypothetical
proportion
mRNA half-life quantile
C
mRNA
protein
All
6185 genes
4085 genes
ρ=-0.2 (p=6x10-55)
ρ=-0.016 (p=0.3)
log2( fold-change )
LoF
1332 genes
1273 genes
ρ=-0.1 (p=1.4x10-4)
ρ=-0.12 (p=1.4x10-5)
Half life (h)
E
mRNA
protein
2000
600
200
Half life (h)
early
mid/early
mid
mid/late
late
constant
no effect

### Suppl. Fig. 7

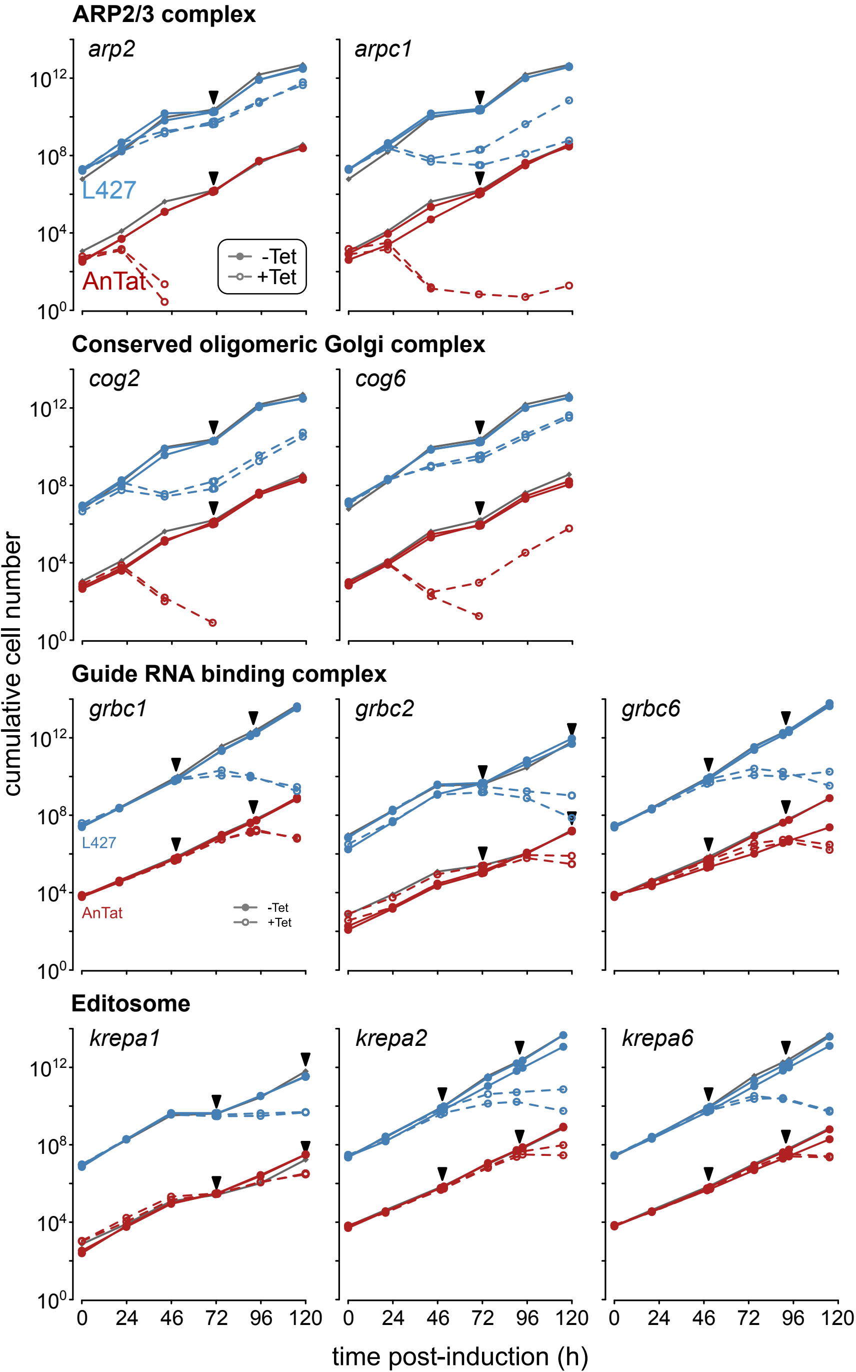
ARP2/3 complex
arp2
arpc1
L427
AnTat
-Tet
+Tet
Conserved oligomeric Golgi complex
cog2
cog6
Guide RNA binding complex
grbc1
grbc2
grbc6
Editosome
krepa1
krepa2
krepa6
cumulative cell number
$10^0$
$10^4$
$10^8$
$10^{12}$
0
24
46
72
96
120
time post-induction (h)

Suppl. Fig. 8

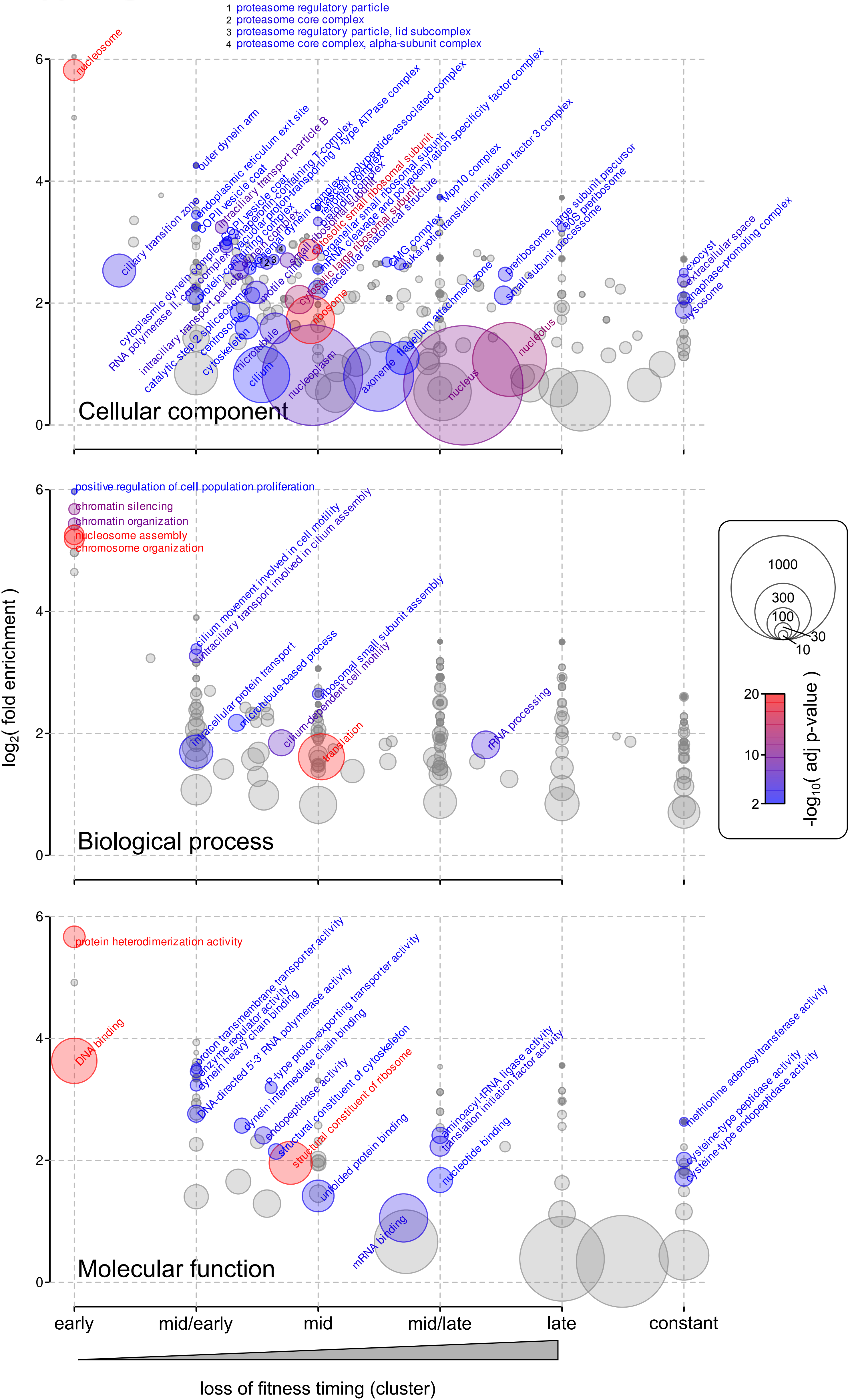
1 proteasome regulatory particle
2 proteasome core complex
3 proteasome regulatory particle, lid subcomplex
4 proteasome core complex, alpha-subunit complex
nucleosome
outer dynein arm
ciliary transition zone
endoplasmic reticulum exit site
COPII vesicle coat
intraciliary transport particle B
chaperonin-containing T-complex
proton-transporting V-type ATPase complex
nascent polypeptide-associated complex
retromer complex
cytosolic small ribosomal subunit
small-subunit processome
organellar small ribosomal subunit
mRNA cleavage and polyadenylation specificity factor complex
cytosolic large ribosomal subunit
intracellular anatomical structure
Mpp10 complex
eukaryotic translation initiation factor 3 complex
MCM complex
preribosome, large subunit precursor
90S preribosome
small-subunit processome
exocyst
extracellular space
anaphase-promoting complex
lysosome
cytoplasmic dynein complex
RNA polymerase II, core complex
protein-containing complex
intraciliary transport particle
catalytic step 2 spliceosome
centrosome
cytoskeleton
microtubule
cilium
motile cilium
ribosome
nucleoplasm
axoneme
flagellum attachment zone
nucleus
nucleolus
Cellular component
positive regulation of cell population proliferation
chromatin silencing
chromatin organization
nucleosome assembly
chromosome organization
cilium movement involved in cell motility
intraciliary transport involved in cilium assembly
intracellular protein transport
microtubule-based process
ribosomal small subunit assembly
cilium-dependent cell motility
translation
rRNA processing
Biological process
protein heterodimerization activity
DNA binding
proton transmembrane transporter activity
enzyme regulator activity
dynein heavy chain binding
DNA-directed 5'-3' RNA polymerase activity
P-type proton-exporting transporter activity
dynein intermediate chain binding
endopeptidase activity
structural constituent of cytoskeleton
structural constituent of ribosome
unfolded protein binding
mRNA binding
aminoacyl-tRNA ligase activity
translation initiation factor activity
nucleotide binding
methionine adenosyltransferase activity
cysteine-type peptidase activity
cysteine-type endopeptidase activity
Molecular function
log2( fold enrichment )
0
2
4
6
early
mid/early
mid
mid/late
late
constant
loss of fitness timing (cluster)
1000
300
100
30
10
20
10
2
-log10( adj p-value )
